# Memory B cell responses to Omicron subvariants after SARS-CoV-2 mRNA breakthrough infection

**DOI:** 10.1101/2022.08.11.503601

**Authors:** Zijun Wang, Pengcheng Zhou, Frauke Muecksch, Alice Cho, Tarek Ben Tanfous, Marie Canis, Leander Witte, Brianna Johnson, Raphael Raspe, Fabian Schmidt, Eva Bednarski, Justin Da Silva, Victor Ramos, Shuai Zong, Martina Turroja, Katrina G. Millard, Kai-Hui Yao, Irina Shimeliovich, Juan Dizon, Anna Kaczynska, Mila Jankovic, Anna Gazumyan, Thiago Y. Oliveira, Marina Caskey, Christian Gaebler, Paul D. Bieniasz, Theodora Hatziioannou, Michel C. Nussenzweig

**Affiliations:** Laboratory of Molecular Immunology, The Rockefeller University, New York, NY 10065, USA; Laboratory of Retrovirology, The Rockefeller University, New York, NY 10065, USA; Howard Hughes Medical Institute

## Abstract

Individuals that receive a 3^rd^ mRNA vaccine dose show enhanced protection against severe COVID19 but little is known about the impact of breakthrough infections on memory responses. Here, we examine the memory antibodies that develop after a 3^rd^ or 4^th^ antigenic exposure by Delta or Omicron BA.1 infection, respectively. A 3^rd^ exposure to antigen by Delta breakthrough increases the number of memory B cells that produce antibodies with comparable potency and breadth to a 3^rd^ mRNA vaccine dose. A 4^th^ antigenic exposure with Omicron BA.1 infection increased variant specific plasma antibody and memory B cell responses. However, the 4^th^ exposure did not increase the overall frequency of memory B cells or their general potency or breadth compared to a 3^rd^ mRNA vaccine dose. In conclusion, a 3^rd^ antigenic exposure by Delta infection elicits strain-specific memory responses and increases in the overall potency and breadth of the memory B cells. In contrast, the effects of a 4^th^ antigenic exposure with Omicron BA.1 is limited to increased strain specific memory with little effect on the potency or breadth of memory B cell antibodies. The results suggest that the effect of strain-specific boosting on memory B cell compartment may be limited.

## Introduction

Severe Acute Respiratory Syndrome Coronavirus (SARS-CoV-2) emerged in late 2019 causing a global pandemic with more than 500 million infections and over 6 million deaths reported to date. Over the course of the pandemic SARS-COV-2 continued to evolve resulting in substantial genetic distance between circulating variants and the initial viral sequence on which vaccines are based. Several of these circulating variants have been designated variants of concern (VoC) and have led to successive waves of infection, most notably by VoCs Alpha (Supasa et al., 2021), Delta (Liu et al., 2021), Omicron (Dejnirattisai et al., 2022).

Higher rates of re-infection and vaccine-breakthrough infection with the Delta and Omicron variants highlighted the potential for immune escape from neutralizing antibody responses resulting in reduced vaccine efficacy against SARS-CoV-2 infection(Cao et al.; Cele et al., 2022; Dejnirattisai et al., 2022; Gaebler et al., 2022; Hachmann et al., 2022; Kuhlmann et al., 2022; Liu et al., 2021). With the emergence of Omicron BA.1 and related lineages infection has surged worldwide, and these new variants account for over 95% of recent COVID-19 cases. To date, BA.2.12.1 variant (a BA.2 lineage) contributes 59% of new cases in the United States, while BA.4 and BA.5 caused a fifth wave of COVID-19 infection in South Africa. Nevertheless, vaccine-elicited immunity continues to provide robust protection against severe disease, even in the face of viral variants(Andrews et al., 2022; Madhi et al., 2022; Wolter et al., 2022; World Health, 2022).

Previous studies have shown that Delta or Omicron breakthrough infection boosts plasma neutralizing activity against both the Wuhan-Hu-1 strain and the infecting variant, which might suggest recall responses of cross-reactive vaccine-induced memory B cells (Kaku et al., 2022; Quandt et al., 2022; Richardson et al., 2022; Seaman et al., 2022; Servellita et al., 2022). However, far less is known about the memory antibody responses after breakthrough infection. Here we report on the development of antibodies produced by memory B cells in a cohort of vaccinated individuals that were subsequently infected with Delta or Omicron.

## Results

Between August 13, 2021, and February 3, 2022, we recruited individuals that had been vaccinated with 2 or 3 doses of an mRNA vaccine and experienced breakthrough infections with Delta (n=24, age range 21 – 63 years, median age=30 years; 67% male, 33% female) or Omicron (n=29, age range 22 – 79 years, median age=33.5 years; 53% male, 47% female) (Table S1, (Gaebler et al., 2022)). Volunteers received either the Moderna (mRNA-1273; n=12), Pfizer-BioNTech (BNT162b2; n=33) or combination (Moderna-Pfizer; n=8) mRNA vaccine (Table S1). Samples from Delta and Omicron BA.1 breakthrough participants were collected a median 26.5 days (range 0-60) or 24 days (range 10-37) after positive test for infection, respectively (Table S1). As a result, Delta breakthrough samples were collected at a median of 5.5 months (range 109-211 days) after 2^nd^ vaccination, and Omicron BA.1 samples were collected at median of 2.4 months (range 26-141 days) after 3^rd^ vaccination (Fig. 1a, see Methods and Table S1). For two participants paired samples were collected shortly after their 3^rd^ vaccine dose and again after Omicron BA.1 breakthrough infection (Table S1).

**Fig. 1:**
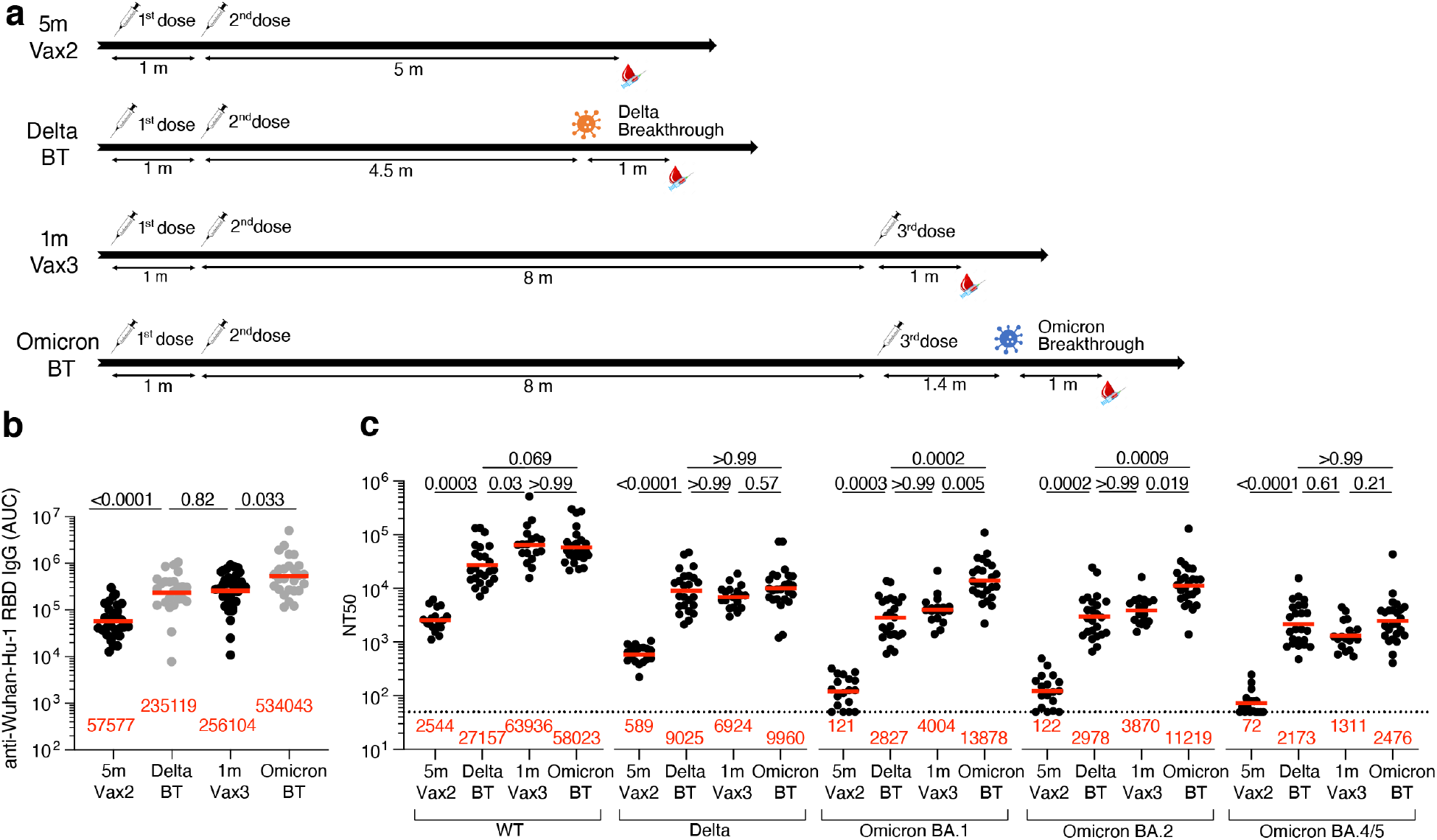
Plasma ELISAs and neutralizing activity. **a**, Diagram shows, blood donation schedules for vaccinated-only individuals 5 months after the 2^nd^ dose (Vax2, top)(Cho et al., 2021), Delta breakthrough infection after Vax2 (Delta BT, 2^nd^ from top), and vaccinated-only individuals 1 month after the 3^rd^ dose (Vax3, 2^nd^ from bottom)(Muecksch et al., 2022) and Omicron breakthrough infection after Vax3 (Omicron BT, bottom). **b**, Graph shows area under the curve (AUG) for plasma IgG antibody binding to wild-type SARS-CoV-2 (WT) RBD after Vax2(Cho et al., 2021), Delta BT for n=24 samples, Vax3(Muecksch et al., 2022) and Omicron BA.1 BT for n=26 samples. **c**, Plasma neutralizing activity against indicated SARS-CoV-2 variants after Vax2(Cho et al., 2021) for n=18 samples, Delta BT for n=24 samples, Vax3(Muecksch et al., 2022) for n=18 samples and Omicron BA.1 BT for n=26 samples. WT and Omicron BA.1 NT_50_ values are derived from two previous reports (Gaebler et al., 2022; Schmidt et al., 2022). See Methods for a list of all substitutions/deletions/insertions in the spike variants. All experiments were performed at least in duplicate. Red bars and values in **a, b**, and **c** represent geometric mean values. Statistical significance in **b** and **c** was determined by two-tailed Kruskal-Wallis test with subsequent Dunn’s multiple comparisons.

## Plasma binding and neutralization

Plasma IgG antibody titers against SARS-CoV-2 Wuhan-Hu-1-(wildtype, WT), or Delta-receptor binding domain (RBD), and Omicron BA.1-Spike were measured by enzyme-linked immunosorbent assays (ELISA) (Wang et al., 2021d). Anti-WT-RBD IgG titers were significantly increased after Delta breakthrough infection in individuals who received 2 doses of mRNA vaccine (Delta BT), compared to vaccinated individuals who did not experience infection (5m-Vax2) (p<0.0001, Vax2 (Cho et al., 2021) vs. Delta, Fig. 1b and Table S1). Similarly, there was a 2-fold increase in geometric mean IgG-binding titers against WT-RBD after Omicron BA.1 breakthrough infection (Omicron BT) in individuals who received 3 doses of mRNA vaccine, compared to vaccinated individuals who were not infected after the 3^rd^ vaccine dose (1m-Vax3) (p=0.033, Vax3 (Muecksch et al., 2022) vs. Omicron, Fig. 1b and Table S1). Individuals who experienced Omicron BA.1 infection exhibited higher anti-Delta-RBD and anti-Omicron BA.1-Spike IgG binding titers than individuals with Delta breakthrough infection or those receiving 3 mRNA vaccine doses (Fig. S1, anti-Delta RBD: p<0.0001, Delta BT vs Omicron BT, p=0.047, Vax3 vs Omicron BT; anti-Omicron BA.1 Spike: p<0.0001, Delta BT vs Omicron BT, p=0.021, Vax3 vs Omicron BT).

Plasma neutralizing activity in 49 participants was measured using HIV-1 pseudotyped with the WT SARS-CoV-2 spike protein (Cho et al., 2021; Wang et al., 2021d) (Fig. 1c and Table S1). Compared to individuals that received 2 mRNA vaccine doses(Cho et al., 2021), Delta breakthrough infection resulted in 11-fold increased geometric mean half-maximal neutralizing titer (NT_50_) (p=0.0003, Vax2 vs. Delta, Fig. 1c). However, the resulting geometric mean NT_50_ was lower than after the 3^rd^ mRNA vaccine dose (p=0.03, Delta vs. Vax3, Fig. 1c). Notably, the NT_50_ against WT after Omicron breakthrough was not significantly different from individuals that received a 3^rd^ vaccine dose (Muecksch et al., 2022) (p>0.99, Vax3 vs. Omicron, Fig. 1c).

Plasma neutralizing activity was also assessed against SARS-CoV-2 Delta, Omicron BA.1, BA.2 and BA.4/5 variants using viruses pseudotyped with appropriate variant spike proteins Delta breakthrough infection resulted in 15-fold increased neutralizing titers against Delta compared to 2-dose vaccinated-only individuals (p<0.0001, Fig. 1c) with resulting titers being comparable to 3-dose vaccinated individuals before and after Omicron breakthrough infection (p>0.99, Fig. 1c). While Delta breakthrough infection also increased neutralizing titers against Omicron BA.1, BA.2, and BA4/5 (p=0.0003, p=0.002 and p<0.0001, respectively, Fig. 1c), the titers were not significantly different from titers observed in 3-dose vaccinated individuals (Muecksch et al., 2022)(p>0.99, p>0.99 and p=0.61, respectively, Fig. 1c). Conversely, Omicron breakthrough infection after 3-dose vaccination resulted in a further 3.5-fold and 2.9-fold increase of Omicron BA.1 and BA.2 neutralizing titers, respectively, when compared to 3-dose vaccinated only individuals (Muecksch et al., 2022)(p=0.005 and p=0.019, respectively. Fig. 1c). Omicron BA.4/5 showed the highest neutralization resistance of all variants tested, resulting in low geometric mean neutralizing titers in plasma samples obtained after the second vaccine dose (NT_50_=72, Fig. 1c). Nevertheless, individuals who had at least 3 antigen exposures (Delta breakthrough, Vax3 and Omicron breakthrough) were able to neutralize Omicron BA.4/5 with NT_50_s of 2173, 1311 and 2476, respectively at the time points assayed.

## Memory B cells

mRNA vaccines elicit memory B cells (MBCs) that can contribute to durable immune protection from serious disease by mediating rapid and anamnestic antibody response (Muecksch et al., 2022; Victora and Nussenzweig, 2022). To better understand the MBC compartment after Delta or Omicron BA.1 breakthrough infection in vaccinated individuals, we enumerated RBD-specific MBCs using Alexa Fluor 647 (AF647)- and phycoerythrin (PE)-labeled WT RBD of the SARS-CoV-2 spike protein by flow cytometry (Fig. 2a, Fig. S2). The number of WT RBD-specific MBCs after Delta breakthrough infection was significantly higher than after the 2^nd^ or 3^rd^ vaccine dose (Delta vs. Vax2, p<0.0001, and Delta vs. Vax3, p=0.011, Fig. 2b). Omicron BA.1 breakthrough infection elicited a 1.7-fold increase in the number of MBCs compared to individuals that received 3 vaccine doses (Vax3 vs. Omicron, p=0.013, Fig. 2b). Consistent with previous reports (Goel et al., 2022; Kaku et al., 2022; Nutalai et al., 2022; Park et al., 2022), flow cytometry showed that a larger fraction of the MBCs developing after the 3^rd^ vaccine dose or Omicron BA.1 breakthrough infection were cross-reactive with WT-, Delta- and Omicron BA.1-RBDs than after Delta breakthrough infection (Fig. 2c). Additional phenotyping indicated that RBD-specific memory B cells elicited by Vax3 or Delta or Omicron BA.1 breakthrough infection showed higher frequencies of IgG than IgM and IgA expression (Fig. S2c-e).

**Fig. 2:**
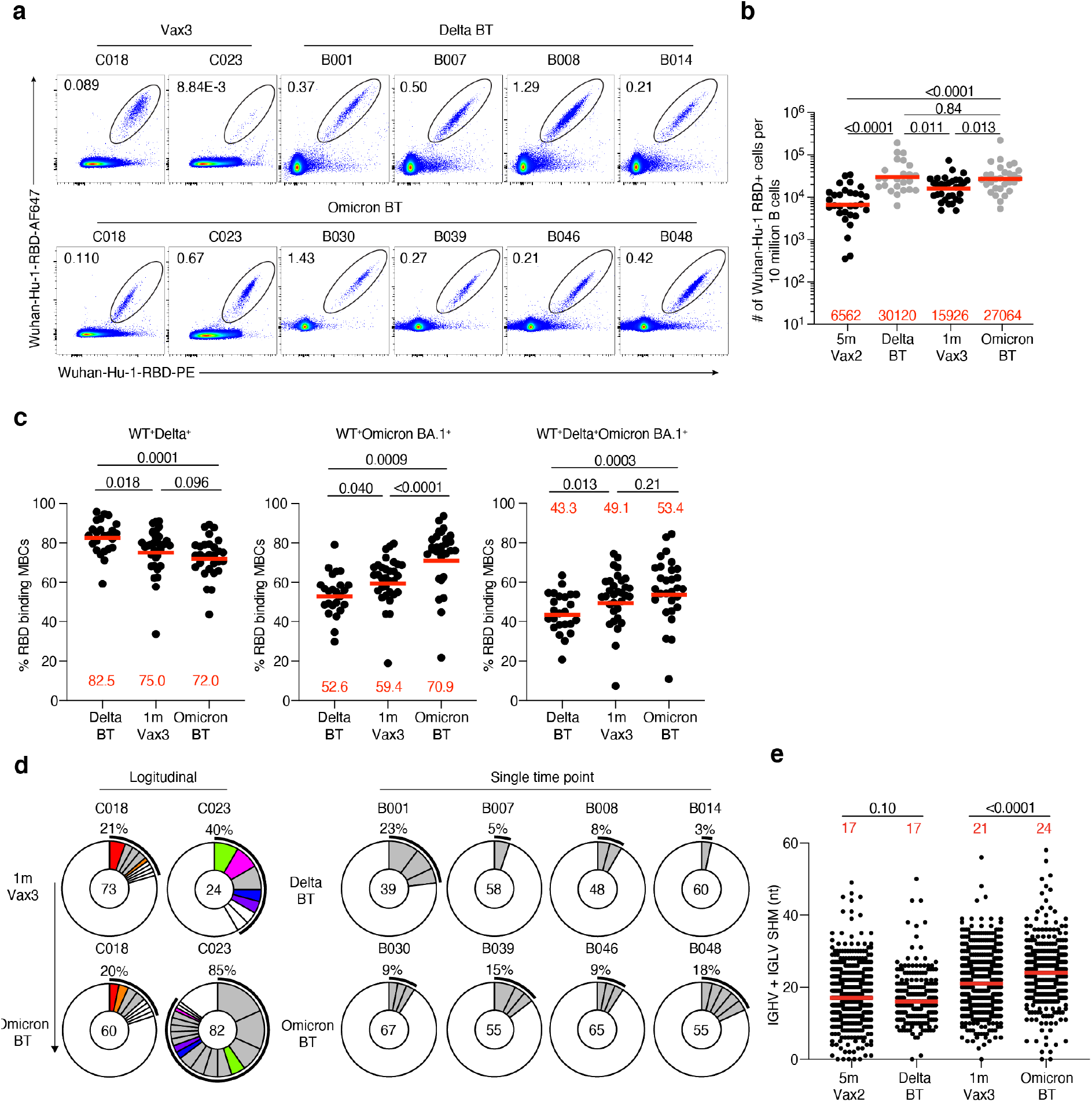
Anti-SARS-CoV-2 RBD memory B cells after breakthrough infection. **a**, Representative flow cytometry plots indicating PE-WT-RBD and AlexaFluor-647-WT-RBD binding memory B cells from 4 individuals after Delta breakthrough infection following Vax2 (Delta BT), 2 individuals 1 month after Vax3, and 6 individuals after Omicron BA.1 breakthrough infection following Vax3 (Omicron BT). The number of WT RBD-specific B cells is indicated in **b**, 5 months after Vax2(Cho et al., 2021), Delta BT (n=24), 1 month after Vax3(Muecksch et al., 2022) and Omicron BT (n=29). **c**, Graphs showing the percentage of WT-, Delta-, and Omicron BA.1-RBD cross-binding B cells determined by flow cytometer in vaccinees (Vax3) and breakthrough individuals (Delta BT) or (Omicron BT) (See also in **Fig. S2b**). **d**, Pie charts show the distribution of IgG antibody sequences obtained from WT-specific memory B cells from: 2 individuals assayed sequentially 1 month after the 3^rd^ mRNA dose (Vax3) an Omicron infection (left); 4 individuals after Delta breakthrough (Delta), and 4 individuals after Omicron breakthrough (Omicron). The number inside the circle indicates the number of sequences analyzed for the individual denoted above the circle. Pie slice size is proportional to the number of clonally related sequences. The black outline and associated numbers indicate the percentage of clonal sequences detected at each time point. Colored slices indicate persisting clones (same IGHV and IGLV genes, with highly similar CDR3s) found at more than one timepoint within the same individual. Grey slices indicate clones unique to the timepoint. White slices indicate sequences isolated only once per time point. **e**, Number of nucleotide somatic hypermutations (SHM) in IGHV + IGLV in WT-RBD-specific sequences after Delta or Omicron breakthrough infection, compared to 5 months after Vax2, and 1 month after Vax3. Red bars and numbers in **b** and **c** represent geometric mean, and in **e**, represent median values. **e**, Statistic analysis in **b**, and **c**, was determined by two-tailed Kruskal-Wallis test with subsequent Dunn’s multiple-comparisons test and in **e** by two-tailed Mann-Whitney test.

To examine the specificity and neutralizing activity of the antibodies produced by MBCs, we purified and sequenced antibody genes in individual WT-RBD-specific B cells from 10 individuals that experienced Delta or Omicron BA.1 breakthrough infection, following the 2^nd^ or 3^rd^ vaccine dose, respectively (Fig. 2d, Fig. S2f, Table S1), including 2 participants for whom paired samples were collected shortly after their 3^rd^ vaccine dose and after subsequent Omicron BA.1 breakthrough infection.

686 paired heavy and light chain anti-RBD antibody sequences were obtained (Fig. 2d, Table S2). Clonally expanded WT-RBD-specific B cells represented 9% of all memory B cells after Delta breakthrough infection and 28% of the repertoire after Omicron BA.1 breakthrough infection (Fig. 2d and Table S2). Similar to mRNA vaccinees (Cho et al., 2021; Muecksch et al., 2022; Wang et al., 2021d), several sets of VH genes including *VH3-30* and *VH3-53* were over-represented in Delta- or Omicron BA.1 breakthrough infection (Fig. S3). In addition, *VH3-49, VH4-38* and *VH1-24* were exclusively over-represented after Delta breakthrough infection (Fig. S3a), while *VH1-69, VH1-58, VH4-61*, and *VH4-38* were specifically over-represented after Omicron BA.1 breakthrough infection (Fig. S3d). These results suggest that Delta and Omicron BA.1 breakthrough infections elicit variant-specific memory antibody responses. While levels of somatic mutation in memory B cells emerging after Delta breakthrough infection were comparable to those after the 2^nd^ vaccine dose, significantly higher numbers of somatic mutations were noted following Omicron BA.1 breakthrough infection compared to the 3^rd^ vaccine dose (p<0.0001) (Fig. 2e, Fig. S4a and b). Moreover, phylogenetic analysis revealed that sequences found after the 3^rd^ vaccine dose and following Omicron BA.1 breakthrough infection were intermingled and similarly distant from their unmutated common ancestors (Fig. S4c).

## Monoclonal antibodies

338 anti-RBD monoclonal antibodies were expressed and tested for binding by ELISA, including 115 antibodies obtained after Delta breakthrough infection (Delta BT), 40 isolated from 2 longitudinal samples after their 3^rd^ vaccine dose in individuals that were subsequently infected (Vax3), and 183 antibodies obtained from 6 individuals after Omicron BA.1 breakthrough infection (Omicron). 85% (n=288) of the antibodies bound to the WT RBD with an EC_50_ of less than 1000 ng/mL (Table S3). The geometric mean ELISA half-maximal concentration (EC_50_) against WT RBD for the monoclonal antibodies obtained from Vax3 was comparable to those found after Delta or Omicron BA.1 breakthrough infections (Fig. 3a). In addition, antibodies isolated after both Delta and Omicron-breakthrough infection showed comparable affinity for WT RBD to antibodies obtained from Vax3 when measured by biolayer interferometry (BLI, Fig. S5a). However, when tested against Delta-RBD antibodies obtained after Delta breakthrough infection showed increased binding compared to those after Vax3. In contrast, there was no statistically significant difference in binding to Omicron BA.1-Spike by Omicron and Vax3 antibodies (Fig. 3a).

**Fig. 3:**
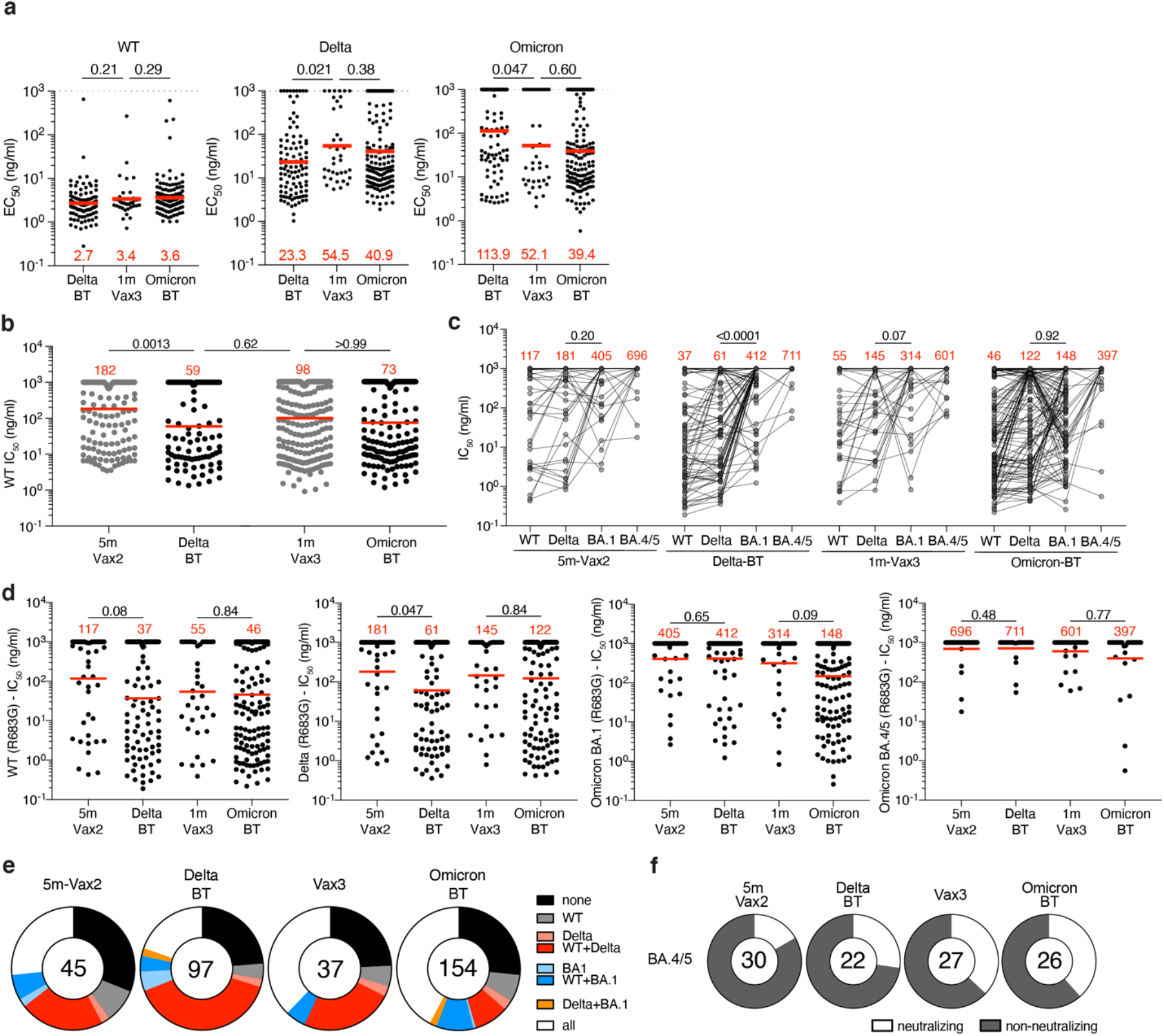
Anti-SARS-CoV-2 RBD monoclonal antibodies. **a**, Graphs show half-maximal effective concentration (EC_50_) of n=342 monoclonal antibodies measured by ELISA against WT-RBD, Delta-RBD and Omicron BA.1-spike protein. Antibodies were obtained memory B cells after Delta breakthrough (Delta BT), after mRNA Vax3, and Omicron breakthrough (Omicron BT). **b**, Graph shows anti-SARS-CoV-2 neutralizing activity of monoclonal antibodies measured by a SARS-CoV-2 pseudotype virus neutralization assay using WT SARS-CoV-2 pseudovirus. IC_50_ values for all antibodies including the 288 reported and tested herein, and 350 previously reported (Cho et al., 2021; Muecksch et al., 2022). **c-d**, Graphs show IC_50_s of monoclonal antibodies against WT, Delta-RBD, and Omicron BA.1 SARS-CoV-2 pseudoviruses. Each dot represents one antibody, where 333 total antibodies were tested including the 288 reported herein, and 45 5m-Vax2 antibodies previously reported(Cho et al., 2021; Muecksch et al., 2022). Red values represent geometric mean values. In addition, 105 antibodies distributed over all four cohorts were also tested against Omicron BA.4/5 psuedovirus. **e**, Ring plots show fraction of neutralizing (IC_50_<1000ng/ml) antibodies against WT, Delta-RBD, and Omicron BA.1 SARS-CoV-2 pseudoviruses, and non-neutralizing (IC_50_>1000 ng/ml) antibodies from each time point. **f**, Ring plots show fraction of mAbs that are neutralizing (IC_50_ 1-1000 ng/mL, white), or non-neutralizing (IC_50_>1000 ng/mL, black) against Omicron BA.4/5. Number in inner circles indicates number of antibodies tested. The deletions/substitutions corresponding to viral variants used in **c-f** were incorporated into a spike protein that also includes the R683G substitution, which disrupts the furin cleavage site and increases particle infectivity. Neutralizing activity against mutant pseudoviruses was compared to a wildtype (WT) SARS-CoV-2 spike sequence (NC_045512), carrying R683G where appropriate. Red bars and values in **a, b, and d**, represent geometric mean values. Statistical significance in **a and b** was determined by two-tailed Kruskal-Wallis test with subsequent Dunn’s multiple comparisons, in c was determined by two-tailed Wilcoxon test and in d was determined by two-tailed Mann-Whitney test.

Anti-RBD antibodies elicited by mRNA vaccination target 4 structurally defined classes of epitopes on the SARS-CoV-2 RBD (Barnes et al., 2020a; Cho et al., 2021; Muecksch et al., 2022; Yuan et al., 2020). To compare the epitopes recognized by anti-RBD memory antibodies elicited by mRNA vaccination (Muecksch et al., 2022) and breakthrough infection, we performed BLI competition experiments. A preformed antibody-RBD-complex was exposed to a second antibody recognizing one of four classes of structurally defined antigenic sites (C105 as Class 1; C144 as Class 2, C135 as Class 3 and C2172 as Class 4 (Barnes et al., 2020a; Muecksch et al., 2022) Fig. S5b). Antibodies obtained after Delta (n=48) or Omicron BA.1 (n=49) breakthrough infection were examined, including 30 of 48 from Delta Breakthrough and 30 of 49 from Omicron BA.1 breakthrough with IC_50_s lower than 1000 ng/mL (neutralizing) against WT (Fig. S5c). In general, there was no significant difference in the distribution of targeted epitopes among antibodies obtained following breakthrough infection as compared to those obtained after mRNA vaccination (Fig. S5c).

All 288 WT RBD-binding antibodies were tested for neutralization in a SARS-CoV-2 pseudotype neutralization assay based on the WT SARS-CoV-2 spike(Robbiani et al., 2020; Schmidt et al., 2020b). For comparison, we used a previously characterized set of antibodies isolated after the 2^nd^ (Cho et al., 2021) or the 3^rd^ vaccine dose (Muecksch et al., 2022). Potency against WT was considerably improved after Delta breakthrough infection compared to the 2^nd^ vaccine dose (Vax2) (IC_50_=182 ng/ml vs IC_50_=50ng/ml, p=0.0013, Fig. 3b) but not compared to the 3^rd^ vaccine dose (IC_50_=98 ng/ml, p=0.62, Fig. 3b). In addition, there was no further improvement of neutralizing activity following Omicron BA.1 breakthrough infection compared to the 3^rd^ dose (IC_50_=73 ng/ml) (p>0.99, Fig. 3b).

To examine whether and how neutralizing breadth evolves in vaccinees after Delta or Omicron BA.1 breakthrough infection, we analyzed the 288 newly expressed antibodies obtained from breakthrough individuals and 45 previously described antibodies obtained from Vax2 individuals (Cho et al., 2021) and measured their neutralizing activity against SARS-CoV-2 pseudoviruses carrying amino acid substitutions found in the Delta-RBD and Omicron BA.1 variant. In addition, 105 randomly selected antibodies from all four groups were tested against an Omicron BA.4/5 pseudovirus (Fig. 3c). Neutralizing potency was generally lower against Omicron BA.1 compared to Delta pseudovirus. However, while antibodies obtained 5 months after the 2^nd^ vaccine dose were not significantly more potent against Delta (IC_50_=181ng/ml) vs. BA.1 pseudovirus (IC_50_=405ng/ml) (p=0.20, Fig. 3c), those obtained after subsequent Delta breakthrough infection neutralize Delta with 6.8-fold increased potency compared to BA.1 (p<0.0001, Fig. 3c). In contrast, the ratio of Delta vs. BA.1 IC_50_ in Vax3 antibodies was only 2.2 (p=0.07, Fig. 3c), while antibodies recovered after subsequent omicron breakthrough neutralized Delta and Omicron with similar potencies IC_50_=122 ng/ml for Delta vs. 148 ng/ml for Omicron (p=0.92, Fig. 3c).

Compared to the 2^nd^ vaccine dose, antibodies from Delta breakthrough infection showed increased potency against Delta pseudovirus (181 vs. 61 ng/ml, p=0.047, Fig. 3d). However, there was no significant improvement of antibody potency against Delta, Omicron BA.1 or Omicron BA.4/5 pseudovirus comparing 5m-Vax2 versus Delta breakthrough antibodies. Moreover, there were only 2 fold differences that did not reach statistical significance when comparing Vax3 and Omicron breakthrough antibodies for Omicron BA.1 and BA.4/5 neutralization (Fig. 3d). Notably, Omicron BA.4/5 showed the highest degree of neutralization resistance for all tested antibody groups (Fig. 3c and d). Neutralizing activity of clonally related antibody pairs from participants C018 and C023 was measured against a panel of SARS-CoV-2 pseudoviruses harboring RBD amino acid substitutions representative of variants including Delta and Omicron BA.1. Most pairs of antibodies obtained from clones persisting between the 3^rd^ dose to the following Omicron BA.1 breakthrough infection showed little improvement in antibody breadth within the analyzed pairs (Table S4).

When comparing the fraction of antibodies showing neutralizing activity against Delta or Delta+WT, or Omicron BA.1 or Omicron BA.1+WT, or all three viruses (WT+Delta+Omicron BA.1), it became apparent that antibodies isolated after 2 vaccine doses and subsequent Delta breakthrough infection show the largest proportion of Delta-neutralizing antibodies. Conversely, antibodies isolated after the 3^rd^ vaccine dose and subsequent Omicron BA.1 breakthrough infection show the largest number of antibodies that neutralized all three pseudoviruses (Fig. 3e, Fig. S5d). Vax3 antibodies and Omicron BA.1 breakthrough antibodies were enriched for those neutralizing BA.4/5 with IC_50_ values of less than 1000 ng/ml with 37% and 38% of all tested antibodies neutralizing BA.4/5, respectively, while only 17% and 27% of Vax2 and Delta breakthrough antibodies, respectively neutralized BA.4/5 (Fig. 3f). Thus, in both cases tested a 3^rd^ exposure to antigen increases memory antibody potency and breadth but a 4^th^ exposure with Omicron BA.1 does little more when it occurs in the time frame measured in this study.

## Discussion

Omicron and its subvariants are reported to be more transmissible than any prior VoC and have spurred a resurgence of new cases worldwide(Mallapaty, 2022). While early reports suggested that Omicron may cause less severe illness, recent studies show variant-specific symptoms but similar virulence (Whitaker et al., 2022), and increased resistance to approved vaccine regimens (Nealon and Cowling, 2022).

We and others have shown that a 3^rd^ mRNA vaccine dose boosts plasma antibody responses to SARS-CoV-2 variants including Omicron BA.1 and increases the number, potency and breadth of the antibodies found in the memory B cell compartment (Goel et al., 2022; Muecksch et al., 2022). Although the antibodies in plasma are generally not sufficient to prevent breakthrough infection boosted individuals are protected against serious disease upon breakthrough infection (Kuhlmann et al., 2022; Nemet et al., 2022). Our findings suggest that a 3^rd^ exposure to antigen in the form of Delta breakthrough infection produces similar effects on the overall size of the memory compartment to a 3^rd^ mRNA vaccine dose, and specifically boosts strain-specific responses. In contrast, while a 4^th^ antigen exposure by infection with Omicron elicits strain-specific memory, it has far more modest effects on the overall potency and breadth of memory B cell antibodies. The data suggest that a variant-specific mRNA vaccine boost will increase plasma neutralizing activity and memory B-cells that are specific to the variant and closely related strains but may not elicit memory B cells with better general potency or breadth than the Wuhan-Hu-1-based mRNA vaccine.

Antigenic variation between viral strains as well as the time interval between antigenic exposures are likely important contributors to the observed differences in immune responses. For example, the antigenic distance between Wuhan-Hu-1 and Delta is shorter than that between Wuhan-hu-1 and Omicron BA.1 or between Delta(Liu et al., 2021) and Omicron BA.1(Dejnirattisai et al., 2022), which could in part explain a more limited antibody response and less cross-reactive MBCs even after a 4th antigen exposure with Omicron BA.1. In addition, we found that Delta breakthrough infection resulted in similar Delta specific antibody responses compared to a 3rd mRNA vaccination or Omicron breakthrough infection. This may be partly due to shorter intervals between exposures in the Delta-BT cohort which is consistent with the notion that the duration after antigen exposure is associated with the continued evolution of the humoral response resulting in greater somatic hypermutation and breadth as well as increased potency (Cho et al., 2021; Gaebler et al., 2021b; Sokal et al., 2021; Wang et al., 2021c).

The data highlights the challenges involved in selecting variant-specific vaccines in the absence of reliable information on the nature of the next emerging variant and suggests that a focus should be on designing vaccines with broader general activity against coronaviruses.

## Supplementary Figure Legends

**Fig. S1:**
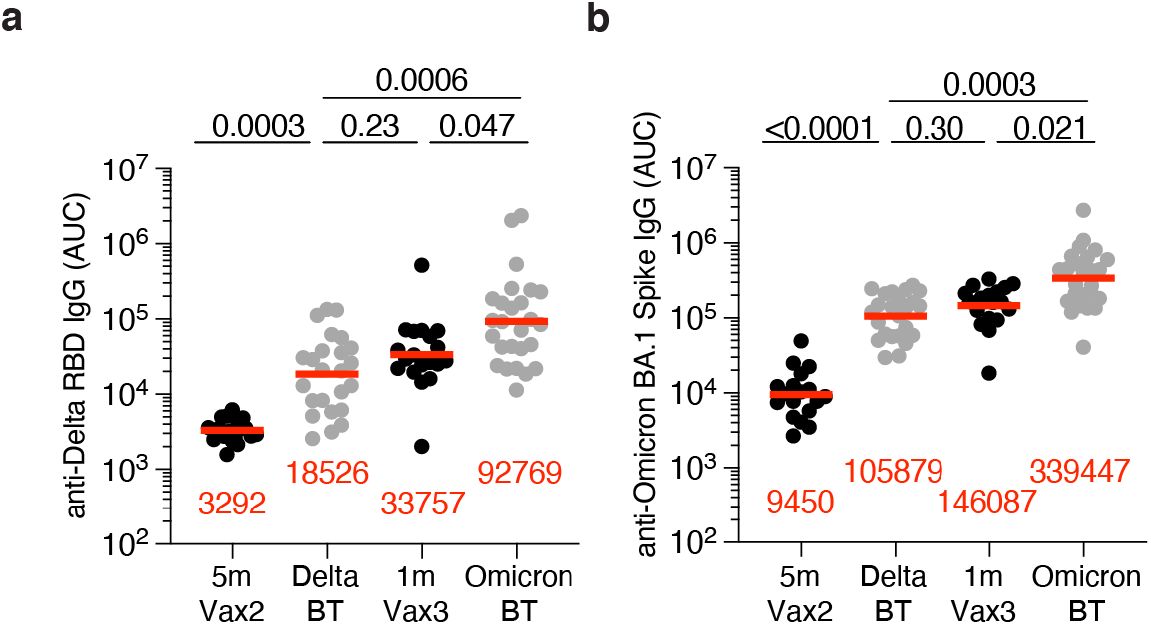
Plasma ELISA. **a-b**, Graph shows area under the curve (AUC) for plasma IgG binding to **a**, SARS-CoV-2 Delta-RBD and **b**, Omicron**-**Spike for vaccinated individuals after Vax2(Cho et al., 2021), Delta breakthrough (Delta BT, n=24), and vaccinated individuals after Vax3 (Muecksch et al., 2022) and Omicron breakthrough infection after Vax3 (Omicron BT, n=26). All experiments were performed at least in duplicate and repeated twice. Red bars and values represent geometric mean values. Statistical significance in **a** and **b** was determined by two-tailed Kruskal-Wallis test with subsequent Dunn’s multiple comparisons.

**Fig. S2:**
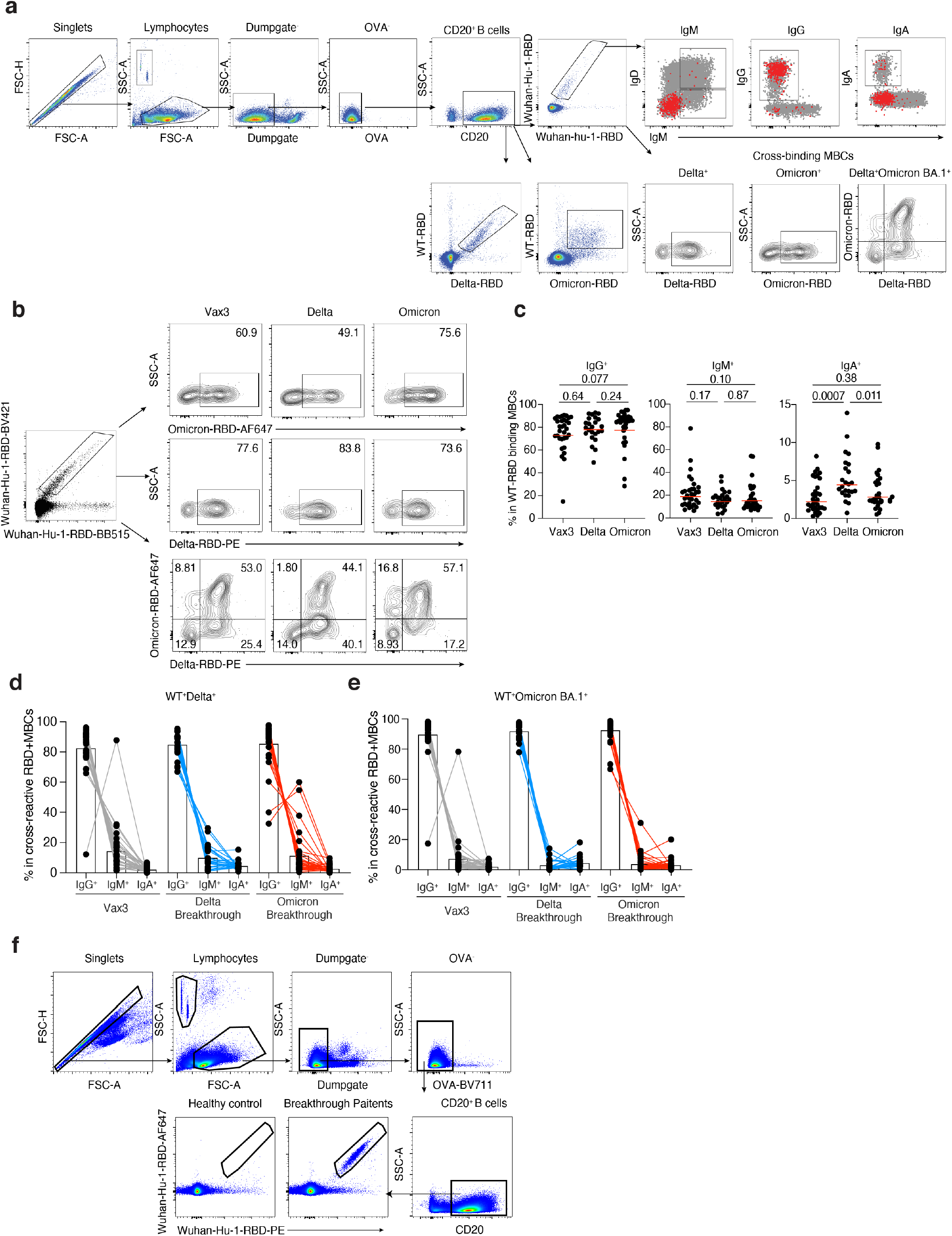
Flow Cytometry. **a-b**, Gating strategy for phenotyping. Gating was on lymphocytes singlets that were CD20^+^ and CD3-CD8-CD16-Ova-. Anti-IgG, IgM, IgA antibodies were used for B cell phenotype analysis. Antigen-specific cells were detected based on binding to WT RBD-PE^+^ and RBD-AF647^+^, or to Delta -RBD and Omicron BA.1-RBD. **c-e**, Graphs show the frequency of IgM, IgG, and IgA isotype expression in **c**, WT RBD+ MBCs, **d**, WT+Delta+ RBD binding MBCs, **e**, WT+Omicron BA.1+ RBD binding MBCs cells. **f**, Gating strategy for single-cell sorting for CD20+ B cells for WT RBD-PE and RBD-AF647. Statistical significance in **c**, was determined by two-tailed Kruskal-Wallis test with subsequent Dunn’s multiple comparisons.

**Fig. S3:**
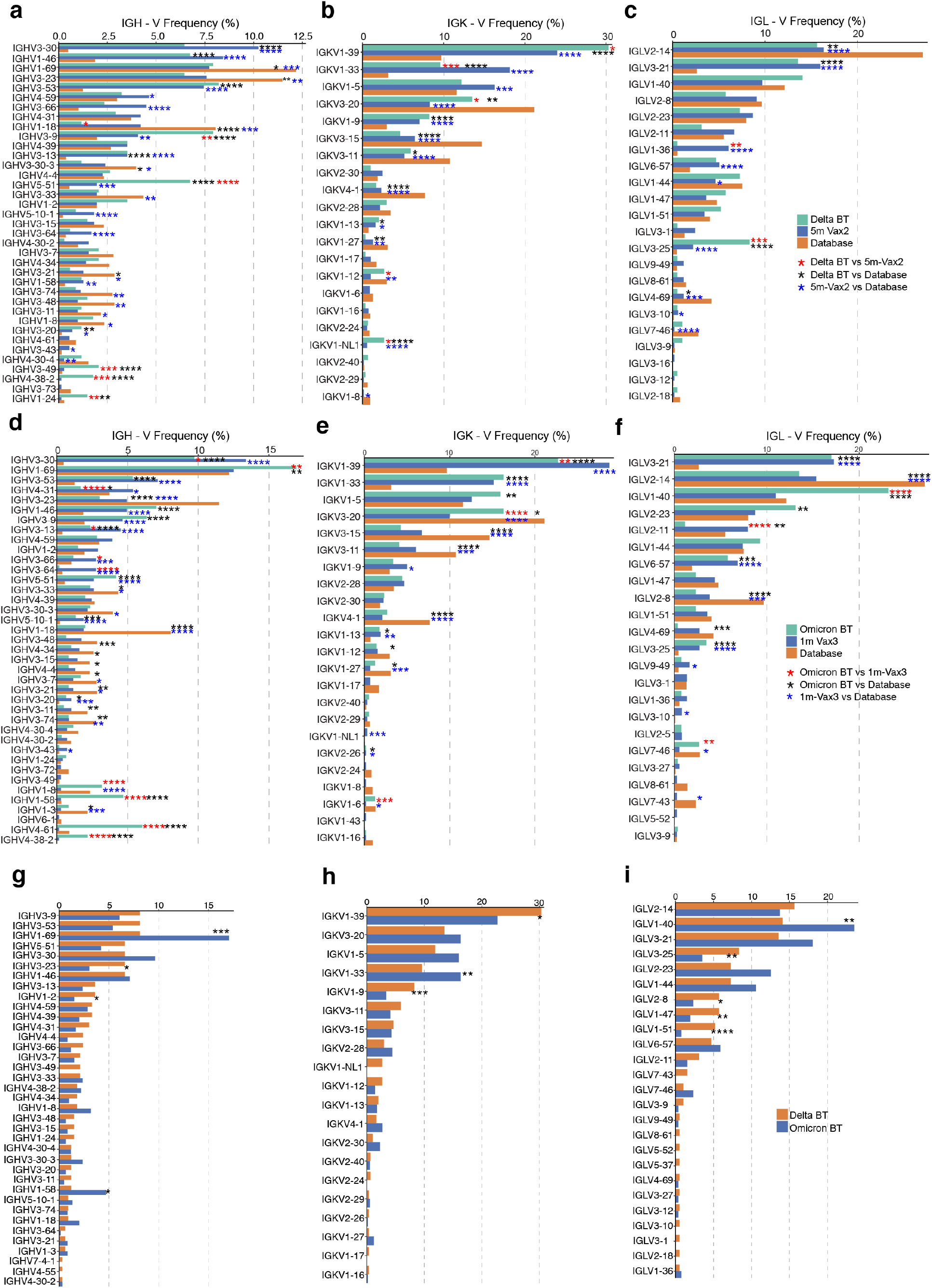
Frequency distribution of human V genes. **a-c**, Comparison of the frequency distribution of human V genes for heavy chain and light chains of anti-RBD antibodies from this study and from a database of shared clonotypes of human B cell receptor generated by Cinque Soto et al (Soto et al., 2019). Graph shows relative abundance of human *IGHV* (left panel), *IGKV* (middle panel) and *IGLV* (right panel) genes in Sequence Read Archive accession SRP010970 (orange), antibodies obtained from Delta breakthrough infection (green), and Vax2 (blue). **d-f**, Same as **a-c**, Graph shows relative abundance of human *IGHV* (left panel), *IGKV* (middle panel) and *IGLV* (right panel) genes in Sequence Read Archive accession SRP010970 (orange), antibodies obtained from Omicron BA.1 breakthrough infection (green), and Vax3 (blue). **g-i**, Graph shows relative abundance of human *IGHV* (left panel), *IGKV* (middle panel) and *IGLV* (right panel) genes of antibodies obtained from Delta breakthrough infection (orange) and from Omicron BA.1 breakthrough infection (blue). Statistical significance was determined by two-sided binomial test. * = p≤0.05, ** = p≤0.01, *** = p≤0.001, **** = p≤0.0001.

**Fig. S4:**
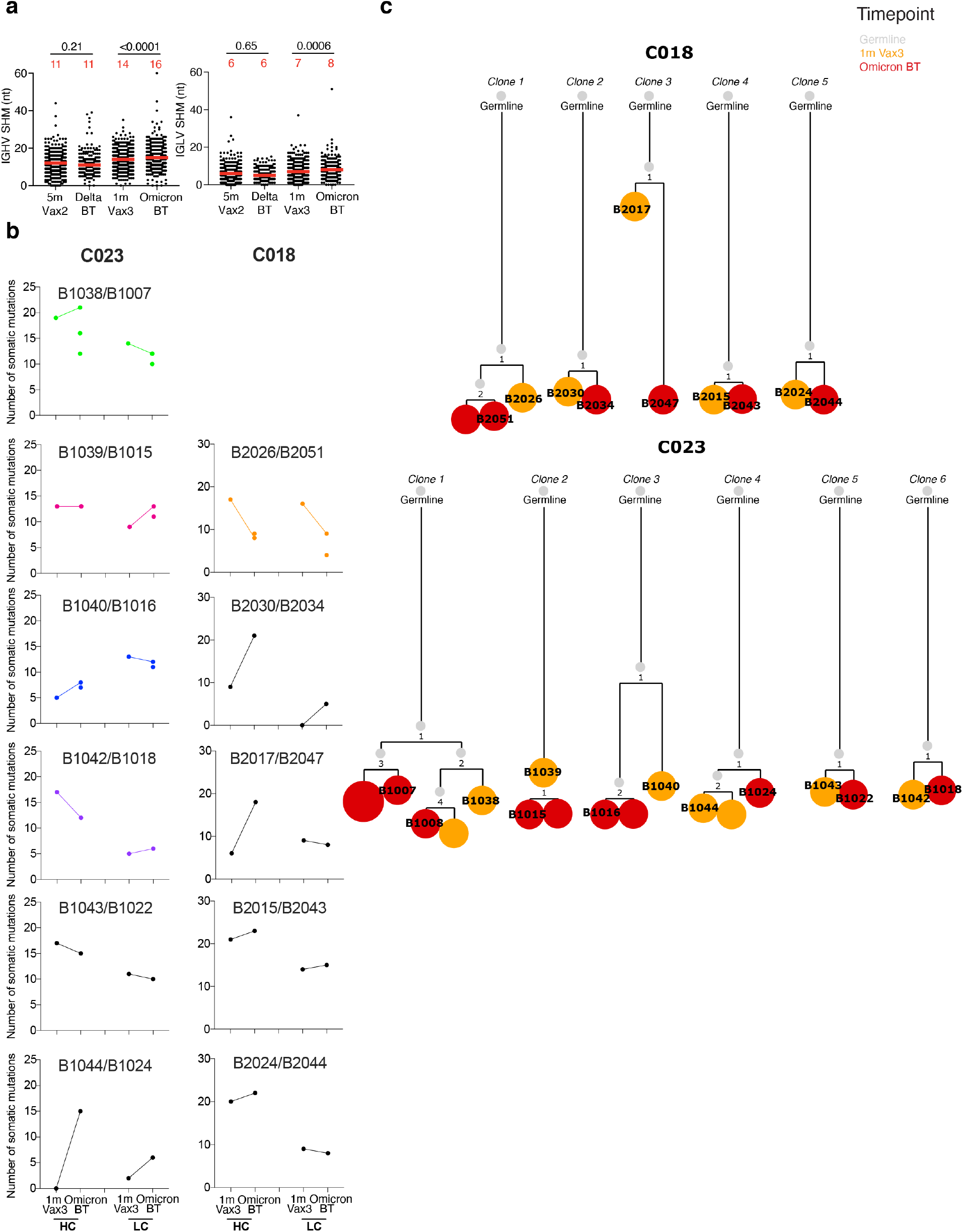
Antibody gene somatic hypermutations analysis and phylogenetic trees. **a**, Number of nucleotide somatic hypermutations (SHM) in *IGHV* and *IGLV* in WT-RBD-specific sequences, separately after Delta or Omicron breakthrough infection, to Vax2 (Cho et al., 2021), and Vax3(Muecksch et al., 2022). Red bars and numbers in **a** represent median value. **b**, The number of somatic nucleotide mutations found in clonally related families found in 1mo after Vax3 and following Omicron breakthrough infection from patients C018 and C023. Color of dot plots match the color of pie slices within the donut plot (Fig 2d), which indicate persisting clones. **c**, the phylogenetic tree graph shows clones from C018 and C023, representing the clonal evolution of RBD-binding memory B cells and derived antibodies obtained from the 3^rd^ mRNA vaccine and the following Omicron BA.1 breakthrough infection.

**Fig. S5:**
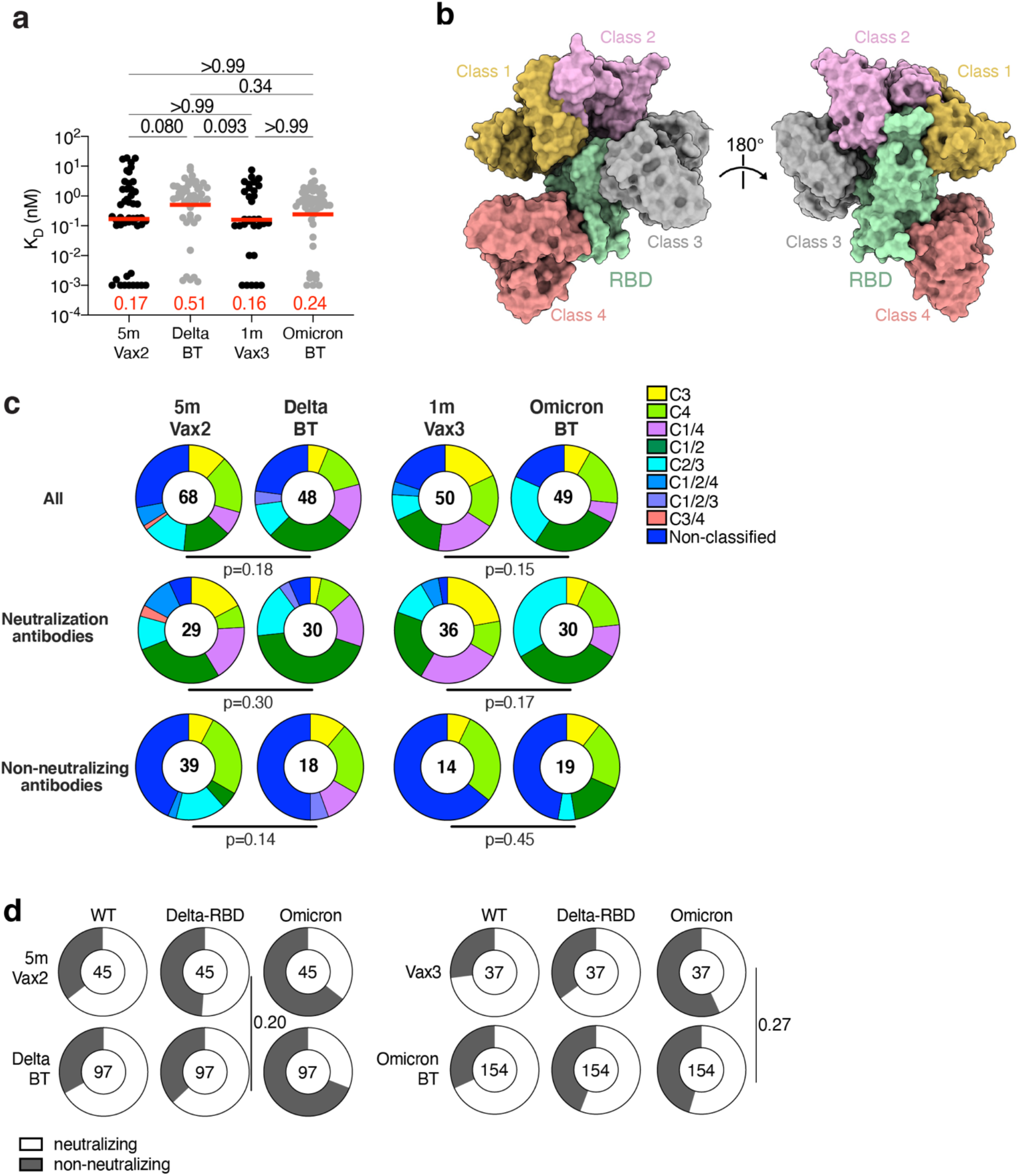
mAb affinity, epitopes and neutralizing breadth. **a**, Graph showing affinity measurements (K_D_s) for WT RBD measured by BLI for antibodies cloned from vaccinated individuals after Delta or Omicron breakthrough infection, compared to Vax2(Cho et al., 2021), and Vax3(Muecksch et al., 2022). **b**, Diagram represents binding poses of antibodies used in BLI competition experiments on the RBD epitope. **c**, Results of epitope mapping performed by competition BLI, comparing mAbs cloned from vaccinated individuals after Delta(n=48) or Omicron BA.1(n=49) breakthrough infection, compared to Vax2(Cho et al., 2021), and Vax3(Muecksch et al., 2022). Pie charts show the distribution of the antibody classes among all RBD-binding antibodies (upper panel), WT neutralizing antibodies only (middle panel) or non-neutralizing antibodies only (lower panel). Red bars represent geometric mean values. Statistical significance was determined by using **a**, by two-tailed Kruskal Wallis test with subsequent Dunn’s multiple comparisons; **c**, two-tailed Chi-square test. **d**, Ring plots show fraction of mAbs in Fig. 3c-e that are neutralizing (IC_50_ 1-1000 ng/mL, white), or non-neutralizing (IC_50_>1000 ng/mL, black) for mutant or variant SARS-CoV-2 pseudovirus indicated across the top at the time point indicated to the left. The number inside the circle indicates the number of antibodies tested. The deletions/substitutions corresponding to viral variants were incorporated into a spike protein that also includes the R683G substitution, which disrupts the furin cleavage site and increases particle infectivity. Neutralizing activity against mutant pseudoviruses was compared to a wildtype (WT) SARS-CoV-2 spike sequence (NC_045512), carrying R683G where appropriate. All experiments were performed at least in duplicate and repeated twice. Statistical significance in **d** was determined by using two-sided Fisher’s exact test.

## Methods

### Study participants

Participants were healthy adults that had been vaccinated with 2 or 3 doses of an mRNA vaccine (mRNA-1273 (Moderna) or BNT162b2 (Pfizer)) and reported breakthrough SARS-CoV-2 infection diagnosed by PCR or antigen testing. Breakthrough infection with delta or omicron variants were deduced based on the prevalent variant circulating in New York City at the time of infection(Gaebler et al., 2022). All participants provided written informed consent before participation in the study and the study was conducted in accordance with Good Clinical Practice. The study was performed in compliance with all relevant ethical regulations and the protocol (DRO-1006) for studies with human participants was approved by the Institutional Review Board of the Rockefeller University. For detailed participant characteristics see Table S1.

### Blood samples processing and storage

Venous blood samples were collected into Heparin and Serum-gel monovette tubes by standard phlebotomy at The Rockefeller University. Peripheral Blood Mononuclear Cells (PBMCs) obtained from samples collected were further purified as previously reported by gradient centrifugation and stored in liquid nitrogen in the presence of Fetal Calf Serum (FCS) and Dimethylsulfoxide (DMSO) (Gaebler et al., 2021a; Robbiani et al., 2020). Heparinized serum and plasma samples were aliquoted and stored at -20°C or less. Prior to experiments, aliquots of plasma samples were heat-inactivated (56°C for 1 hour) and then stored at 4°C.

### ELISAs

Enzyme-Linked Immunosorbent Assays (ELISAs)(Amanat et al., 2020; Grifoni. et al., 2020) were performed to evaluate antibodies binding to SARS-CoV-2 wild-type (Wuhan-Hu-1) RBD, and variants of concern Delta (B.1.617.2) RBD, and Omicron (BA.1) spike protein by coating of high-binding 96-half-well plates (Corning 3690) with 50 μl per well of a 1μg/ml indicated protein solution in Phosphate-buffered Saline (PBS) overnight at 4°C. Plates were washed 6 times with washing buffer (1× PBS with 0.05% Tween-20 (Sigma-Aldrich)) and incubated with 170 μl per well blocking buffer (1× PBS with 2% BSA and 0.05% Tween-20 (Sigma)) for 1 hour at room temperature. Immediately after blocking, plasma samples or monoclonal antibodies were added in PBS and incubated for 1 hour at room temperature. Plasma samples were assayed at a 1:66 starting dilution and 10 additional 3-fold serial dilutions.

10 μg/ml starting concentration was used to test monoclonal antibodies followed by 10 additional 4-fold serial dilutions. Plates were washed 6 times with washing buffer and then incubated with anti-human IgG secondary antibody conjugated to horseradish peroxidase (HRP) (Jackson Immuno Research 109-036-088 109-035-129 and Sigma A0295) in blocking buffer at a 1:5,000 dilution. Plates were developed by addition of the HRP substrate, 3,3’,5,5’-Tetramethylbenzidine (TMB) (ThermoFisher) for 10 minutes (plasma samples and monoclonal antibodies). 50 μl of 1 M H_2_SO_4_ was used to stop the reaction and absorbance was measured at 450 nm with an ELISA microplate reader (FluoStar Omega, BMG Labtech) with Omega and Omega MARS software for analysis. A positive control (For anti-RBD ELISA, plasma from participant COV72, diluted 66.6-fold and ten additional threefold serial dilutions in PBS; for anti-Omicron spike ELISA, plasma from B039 was used as a control) was added to every assay plate for normalization for plasma samples. The average of its signal was used for normalization of all the other values on the same plate with Excel software before calculating the area under the curve using Prism V9.1(GraphPad).

Negative controls of pre-pandemic plasma samples from healthy donors were used for validation (for more details please see(Robbiani et al., 2020)). For monoclonal antibodies, the ELISA half-maximal concentration (EC_50_) was determined using four-parameter nonlinear regression (GraphPad Prism V9.1). EC_50_s above 1000 ng/mL were considered non-binders.

### Proteins

The mammalian expression vector encoding the Receptor Binding-Domain (RBD) of SARS-CoV-2 (GenBank MN985325.1; Spike (S) protein residues 319-539) was previously described(Barnes et al., 2020b).

### SARS-CoV-2 pseudotyped reporter virus

A panel of plasmids expressing RBD-mutant SARS-CoV-2 spike proteins in the context of pSARS-CoV-2-S_Δ19_ has been described(Cho et al., 2021; Muecksch et al., 2021; Wang et al., 2021d; Weisblum et al., 2020). Variant pseudoviruses resembling SARS-CoV-2 variants Delta (B.1.617.2) and Omicron BA.1 (B.1.1.529) have been described before(Cho et al., 2021; Schmidt et al., 2022; Wang et al., 2021b) and were generated by introduction of substitutions using synthetic gene fragments (IDT) or overlap extension PCR mediated mutagenesis and Gibson assembly. Specifically, the variant-specific deletions and substitutions introduced were: Delta: T19R, Δ156-158, L452R, T478K, D614G, P681R, D950N Omicron BA.1: A67V, Δ69-70, T95I, G142D, Δ143-145, Δ211, L212I, ins214EPE, G339D, S371L, S373P, S375F, K417N, N440K, G446S, S477N, T478K, E484A, Q493K, G496S, Q498R, N501Y, Y505H, T547K, D614G, H655Y, H679K, P681H, N764K, D796Y, N856K, Q954H, N969H, N969K, L981F Omicron BA.2: T19I, L24S, del25-27, G142D, V213G, G339D, S371F, S373P, S375F, T376A, D405N, R408S, K417N, N440K, S477N, T478K, E484A, Q493R, Q498R, N501Y, Y505H, D614G, H655Y, N679K, P681H, N764K, D796Y, Q954H, N969K Omicron BA.4/5: T19I, L24S, del25-27, del69-70, G142D, V213G, G339D, S371F, S373P, S375F, T376A, D405N, R408S, K417N, N440K, L452R, S477N, T478K, E484A, F486V, Q498R, N501Y, Y505H, D614G, H655Y, N679K, P681H, N764K, D796Y, Q954H, N969K Deletions/substitutions corresponding to variants of concern listed above were incorporated into a spike protein that also includes the R683G substitution, which disrupts the furin cleavage site and increases particle infectivity. Neutralizing activity against mutant pseudoviruses were compared to a wildtype (WT) SARS-CoV-2 spike sequence (NC_045512), carrying R683G where appropriate.

SARS-CoV-2 pseudotyped particles were generated as previously described(Robbiani et al., 2020; Schmidt et al., 2020a). Briefly, 293T (CRL-11268) cells were obtained from ATCC, and the cells were transfected with pNL4-3ΔEnv-nanoluc and pSARS-CoV-2-S_Δ19_, particles were harvested 48 hours post-transfection, filtered and stored at -80°C.

### Pseudotyped virus neutralization assay

Pre-pandemic negative control plasma from healthy donors, plasma from individuals who received mRNA vaccines and had Delta or Omicron BA.1 breakthrough infection, or monoclonal antibodies were five-fold serially diluted and incubated with SARS-CoV-2 pseudotyped virus for 1 hour at 37 °C. The mixture was subsequently incubated with 293T_Ace2_ cells(Robbiani et al., 2020) (for all WT neutralization assays) or HT1080/Ace2 cl14 cells (for all variant neutralization assays) for 48 hours after which cells were washed with PBS and lysedwith Luciferase Cell Culture Lysis 5× reagent (Promega). Nanoluc Luciferase activity in lysateswas measured using the Nano-Glo Luciferase Assay System (Promega) with the ClarioStar Microplate Multimode Reader (BMG). The relative luminescence units were normalized to those derived from cells infected with SARS-CoV-2 pseudotyped virus(Wang et al., 2021d) in the absence of plasma or monoclonal antibodies. The half-maximal neutralization titers for plasma (NT_50_) or half-maximal and 90% inhibitory concentrations for monoclonal antibodies (IC_50_ and IC_90_) were determined using four-parameter nonlinear regression (least squares regression method without weighting; constraints: top=1, bottom=0) (GraphPad Prism).

### Biotinylation of viral protein for use in flow cytometry

Purified and Avi-tagged SARS-CoV-2 WT and Delta RBD was biotinylated using the Biotin-Protein Ligase-BIRA kit according to manufacturer’s instructions (Avidity) as described before(Robbiani et al., 2020). Ovalbumin (Sigma, A5503-1G) was biotinylated using the EZ-Link Sulfo-NHS-LC-Biotinylation kit according to the manufacturer’s instructions (Thermo Scientific). Biotinylated ovalbumin was conjugated to streptavidin-BV711 for single-cell sorts (BD biosciences, 563262) or to streptavidin-BB515 for phenotyping panel (BD, 564453). WT RBD was conjugated to streptavidin-PE (BD Biosciences, 554061) and streptavidin-AF647 (Biolegend, 405237) for single-cell sorts, or streptavidin-BV421 (Biolegend, 405225) and streptavidin-BV711 (BD biosciences, 563262) for phenotyping. Delta RBD was conjugated to streptavidin-PE (BD Biosciences, 554061) and Omicron BA.1 RBD (ACROBiosystems, SPD-C82E4) was conjugated to streptavidin-AF647 (Biolegend, 405237).

### Flow cytometry and single cell sorting

Single-cell sorting by flow cytometry was described previously(Robbiani et al., 2020). Simply, peripheral blood mononuclear cells (PBMC) were enriched for B cells by negative selection using a pan-B-cell isolation kit according to the manufacturer’s instructions (Miltenyi Biotec, 130-101-638). The enriched B cells were incubated in Flourescence-Activated Cell-sorting (FACS) buffer (1 x PBS, 2% FCS, 1 mM ethylenediaminetetraacetic acid (EDTA)) with the following anti-human antibodies (all at 1:200 dilution): anti-CD20-PECy7 (BD Biosciences, 335793), anti-CD3-APC-eFluro 780 (Invitrogen, 47-0037-41), anti-CD8-APC-eFluor 780 (Invitrogen, 47-0086-42), anti-CD16-APC-eFluor 780 (Invitrogen, 47-0168-41), anti-CD14-APC-eFluor 780 (Invitrogen, 47-0149-42), as well as Zombie NIR (BioLegend, 423105) and fluorophore-labeled RBD and ovalbumin (Ova) for 30 min on ice. Single CD3^-^CD8^-^CD14^-^CD16^-^CD20^+^Ova^—^WT RBD-PE^+^-WT RBD-AF647^+^ B cells were sorted into individual wells of 96-well plates containing 4 μl of lysis buffer (0.5 x PBS, 10 mM Dithiothreitol (DTT), 3,000 units/ml RNasin Ribonuclease Inhibitors (Promega, N2615) per well using a FACS Aria III and FACSDiva software (Becton Dickinson) for acquisition and FlowJo for analysis. The sorted cells were frozen on dry ice, and then stored at -80 °C or immediately used for subsequent RNA reverse transcription. For B cell phenotype analysis, in addition to above antibodies, B cells were also stained with following anti-human antibodies (all at 1:200 dilution): anti-IgD-BV650 (BD, 740594), anti-CD27-BV786 (BD biosciences, 563327), anti-CD19-BV605 (Biolegend, 302244), anti-CD71-PerCP-Cy5.5 (Biolegend, 334114), anti-IgG-PECF594 (BD, 562538), anti-IgM-AF700 (Biolegend, 314538), anti-IgA-Viogreen (Miltenyi Biotec, 130-113-481).

### Antibody sequencing, cloning and expression

Antibodies were identified and sequenced as described previously (Robbiani et al., 2020; Wang et al., 2021a). In brief, RNA from single cells was reverse transcribed (SuperScript III Reverse Transcriptase, Invitrogen, 18080-044), and the cDNA was stored at –20 °C or used for subsequent amplification of the variable *IGH, IGL* and *IGK* genes by nested PCR and Sanger sequencing. Sequence analysis was performed using MacVector. Amplicons from the first PCR reaction were used as templates for sequence- and ligation-independent cloning into antibody expression vectors. Recombinant monoclonal antibodies were produced and purified as previously described(Robbiani et al., 2020).

### Biolayer interferometry

Biolayer interferometry assays were performed as previously described(Robbiani et al., 2020). In brief, we used the Octet Red instrument (ForteBio) at 30 °C with shaking at 1,000 r.p.m. Epitope binding assays were performed with protein A biosensor (ForteBio 18-5010), following the manufacturer’s protocol “classical sandwich assay” as follows: (1) Sensor check: sensors immersed 30 sec in buffer alone (buffer ForteBio 18-1105), (2) Capture 1^st^ Ab: sensors immersed 10 min with Ab1 at 10 μg/mL, (3) Baseline: sensors immersed 30 sec in buffer alone, (4) Blocking: sensors immersed 5 min with IgG isotype control at 10 μg/mL. (5) Baseline: sensors immersed 30 sec in buffer alone, (6) Antigen association: sensors immersed 5 min with RBD at 10 μg/mL. (7) Baseline: sensors immersed 30 sec in buffer alone. (8) Association Ab2: sensors immersed 5 min with Ab2 at 10 μg/mL. Curve fitting was performed using the Fortebio Octet Data analysis software (ForteBio). Affinity measurement of anti-SARS-CoV-2 IgGs binding were corrected by subtracting the signal obtained from traces performed with IgGs in the absence of WT RBD. The kinetic analysis using protein A biosensor (as above) was performed as follows: (1) baseline: 60sec immersion in buffer. (2) loading: 200sec immersion in a solution with IgGs 10 μg/ml. (3) baseline: 200sec immersion in buffer. (4) Association: 300sec immersion in solution with WT RBD at 20, 10 or 5 μg/ml (5) dissociation: 600sec immersion in buffer. Curve fitting was performed using a fast 1:1 binding model and the Data analysis software (ForteBio). Mean *K*_D_ values were determined by averaging all binding curves that matched the theoretical fit with an R^2^ value ≥ 0.8.

### Computational analyses of antibody sequences

Antibody sequences were trimmed based on quality and annotated using Igblastn v.1.14. with IMGT domain delineation system. Annotation was performed systematically using Change-O toolkit v.0.4.540 (Gupta et al., 2015). Clonality of heavy and light chain was determined using DefineClones.py implemented by Change-O v0.4.5 (Gupta et al., 2015). The script calculates the Hamming distance between each sequence in the data set and its nearest neighbor. Distances are subsequently normalized and to account for differences in junction sequence length, and clonality is determined based on a cut-off threshold of 0.15. Heavy and light chains derived from the same cell were subsequently paired, and clonotypes were assigned based on their V and J genes using in-house R and Perl scripts. All scripts and the data used to process antibody sequences are publicly available on GitHub (https://github.com/stratust/igpipeline/tree/igpipeline2_timepoint_v2).

The frequency distributions of human V genes in anti-SARS-CoV-2 antibodies from this study was compared to 131,284,220 IgH and IgL sequences generated by(Soto et al., 2019) and downloaded from cAb-Rep(Guo et al., 2019), a database of human shared BCR clonotypes available at https://cab-rep.c2b2.columbia.edu/. We selected the IgH and IgL sequences from the database that are partially coded by the same V genes and counted them according to the constant region. The frequencies shown in Fig. S3 are relative to the source and isotype analyzed. We used the two-sided binomial test to check whether the number of sequences belonging to a specific *IGHV* or *IGLV* gene in the repertoire is different according to the frequency of the same IgV gene in the database. Adjusted p-values were calculated using the false discovery rate (FDR) correction. Significant differences are denoted with stars.

Nucleotide somatic hypermutation and Complementarity-Determining Region (CDR3) length were determined using in-house R and Perl scripts. For somatic hypermutations, *IGHV* and *IGLV* nucleotide sequences were aligned against their closest germlines using Igblastn and the number of differences was considered to correspond to nucleotide mutations. The average number of mutations for V genes was calculated by dividing the sum of all nucleotide mutations across all participants by the number of sequences used for the analysis. GCTree (https://github.com/matsengrp/gctree) (DeWitt et al., 2018) was further used to perform the phylogenetic trees construction. Each node represents a unique IgH and IgL combination and the size of each node is proportional to the number of identical sequences. The numbered nodes represent the unobserved ancestral genotypes between the germline sequence and the sequences on the downstream branch.

## Data presentation

Figures were arranged in Adobe Illustrator 2022.

## Data availability statement

Data are provided in Tables S1-4. The raw sequencing data and computer scripts associated with Figure 2 have been deposited at Github (https://github.com/stratust/igpipeline/tree/igpipeline2_timepoint_v2). This study also uses data from “A Public Database of Memory and Naive B-Cell Receptor Sequences” (https://doi.org/10.5061/dryad.35ks2), Protein Data Bank (6VYB and 6NB6), cAb-Rep (https://cab-rep.c2b2.columbia.edu/), the Sequence Read Archive (accession SRP010970), and from “High frequency of shared clonotypes in human B cell receptor repertoires” (https://doi.org/10.1038/s41586-019-0934-8).

## Code availability statement

Computer code to process the antibody sequences is available at GitHub (https://github.com/stratust/igpipeline/tree/igpipeline2_time-point_v2).

## Notes

### Competing Interest Statement

The Rockefeller University has filed a provisional patent application in connection with this work on which M.C.N. is an inventor (US patent 63/021,387).

## Reference

Amanat, F., D. Stadlbauer, S. Strohmeier, T.H. Nguyen, V. Chromikova, M. McMahon, K. Jiang, G.A. Arunkumar, D. Jurczyszak, and J. Polanco. 2020. A serological assay to detect SARS-CoV-2 seroconversion in humans. Nature medicine 26:1033–1036.

Andrews, N., J. Stowe, F. Kirsebom, S. Toffa, T. Rickeard, E. Gallagher, C. Gower, M. Kall, N. Groves, A.M. O’Connell, D. Simons, P.B. Blomquist, A. Zaidi, S. Nash, N. Iwani Binti Abdul Aziz, S. Thelwall, G. Dabrera, R. Myers, G. Amirthalingam, S. Gharbia, J.C. Barrett, R. Elson, S.N. Ladhani, N. Ferguson, M. Zambon, C.N.J. Campbell, K. Brown, S. Hopkins, M. Chand, M. Ramsay, and J. Lopez Bernal. 2022. Covid-19 Vaccine Effectiveness against the Omicron (B.1.1.529) Variant. N Engl J Med 386:1532–1546.

Barnes, C.O., C.A. Jette, M.E. Abernathy, K.-M.A. Dam, S.R. Esswein, H.B. Gristick, A.G. Malyutin, N.G. Sharaf, K.E. Huey-Tubman, and Y.E. Lee. 2020a. SARS-CoV-2 neutralizing antibody structures inform therapeutic strategies. Nature 588:682–687.

Barnes, C.O., A.P. West Jr, K.E. Huey-Tubman, M.A. Hoffmann, N.G. Sharaf, P.R. Hoffman, N. Koranda, H.B. Gristick, C. Gaebler, and F. Muecksch. 2020b. Structures of human antibodies bound to SARS-CoV-2 spike reveal common epitopes and recurrent features of antibodies. Cell 182:828-842.e816.

Cao, Y., A. Yisimayi, F. Jian, W. Song, T. Xiao, L. Wang, S. Du, J. Wang, Q. Li, X. Chen, P. Wang, Z. Zhang, P. Liu, R. An, X. Hao, Y. Wang, J. Wang, R. Feng, H. Sun, L. Zhao, W. Zhang, D. Zhao, J. Zheng, L. Yu, C. Li, N. Zhang, R. Wang, X. Niu, S. Yang, X. Song, L. Zheng, Z. Li, Q. Gu, F. Shao, W. Huang, R. Jin, Z. Shen, Y. Wang, X. Wang, J. Xiao, and X.S. Xie. - BA.2.12.1, BA.4 and BA.5 escape antibodies elicited by Omicron infection. - 2022.2004.2030.489997.

Cele, S., L. Jackson, D.S. Khoury, K. Khan, T. Moyo-Gwete, H. Tegally, J.E. San, D. Cromer, C. Scheepers, D.G. Amoako, F. Karim, M. Bernstein, G. Lustig, D. Archary, M. Smith, Y. Ganga, Z. Jule, K. Reedoy, S.H. Hwa, J. Giandhari, J.M. Blackburn, B.I. Gosnell, S.S. Abdool Karim, W. Hanekom, S.A. Ngs, C.-K. Team, A. von Gottberg, J.N. Bhiman, R.J. Lessells, M.S. Moosa, M.P. Davenport, T. de Oliveira, P.L. Moore, and A. Sigal. 2022. Omicron extensively but incompletely escapes Pfizer BNT162b2 neutralization. Nature 602:654–656.

Cho, A., F. Muecksch, D. Schaefer-Babajew, Z. Wang, S. Finkin, C. Gaebler, V. Ramos, M. Cipolla, P. Mendoza, M. Agudelo, E. Bednarski, J. DaSilva, I. Shimeliovich, J. Dizon, M. Daga, K.G. Millard, M. Turroja, F. Schmidt, F. Zhang, T.B. Tanfous, M. Jankovic, T.Y. Oliveria, A. Gazumyan, M. Caskey, P.D. Bieniasz, T. Hatziioannou, and M.C. Nussenzweig. 2021. Anti-SARS-CoV-2 receptor-binding domain antibody evolution after mRNA vaccination. Nature 600:517–522.

Dejnirattisai, W., J. Huo, D. Zhou, J. Zahradnik, P. Supasa, C. Liu, H.M.E. Duyvesteyn, H.M. Ginn, A.J. Mentzer, A. Tuekprakhon, R. Nutalai, B. Wang, A. Dijokaite, S. Khan, O. Avinoam, M. Bahar, D. Skelly, S. Adele, S.A. Johnson, A. Amini, T.G. Ritter, C. Mason, C. Dold, D. Pan, S. Assadi, A. Bellass, N. Omo-Dare, D. Koeckerling, A. Flaxman, D. Jenkin, P.K. Aley, M. Voysey, S.A. Costa Clemens, F.G. Naveca, V. Nascimento, F. Nascimento, C. Fernandes da Costa, P.C. Resende, A. Pauvolid-Correa, M.M. Siqueira, V. Baillie, N. Serafin, G. Kwatra, K. Da Silva, S.A. Madhi, M.C. Nunes, T. Malik, P.J.M. Openshaw, J.K. Baillie, M.G. Semple, A.R. Townsend, K.A. Huang, T.K. Tan, M.W. Carroll, P. Klenerman, E. Barnes, S.J. Dunachie, B. Constantinides, H. Webster, D. Crook, A.J. Pollard, T. Lambe, O. Consortium, I.C. Consortium, N.G. Paterson, M.A. Williams, D.R. Hall, E.E. Fry, J. Mongkolsapaya, J. Ren, G. Schreiber, D.I. Stuart, and G.R. Screaton. 2022. SARS-CoV-2 Omicron-B.1.1.529 leads to widespread escape from neutralizing antibody responses. Cell 185:467–484 e415.

DeWitt, W.S., 3rd, L. Mesin, G.D. Victora, V.N. Minin, and F.A.t. Matsen. 2018. Using Genotype Abundance to Improve Phylogenetic Inference. Mol Biol Evol 35:1253–1265.

Gaebler, C., J. DaSilva, E. Bednarski, F. Muecksch, F. Schmidt, Y. Weisblum, K.G. Millard, M. Turroja, A. Cho, Z. Wang, M. Caskey, M.C. Nussenzweig, P.D. Bieniasz, and T. Hatziioannou. 2022. SARS-CoV-2 neutralization after mRNA vaccination and variant breakthrough infection. Open Forum Infectious Diseases

Gaebler, C., Z. Wang, J.C. Lorenzi, F. Muecksch, S. Finkin, M. Tokuyama, A. Cho, M. Jankovic, D. Schaefer-Babajew, and T.Y. Oliveira. 2021a. Evolution of antibody immunity to SARS-CoV-2. Nature 591:639–644.

Gaebler, C., Z. Wang, J.C.C. Lorenzi, F. Muecksch, S. Finkin, M. Tokuyama, A. Cho, M. Jankovic, D. Schaefer-Babajew, T.Y. Oliveira, M. Cipolla, C. Viant, C.O. Barnes, Y. Bram, G. Breton, T. Hagglof, P. Mendoza, A. Hurley, M. Turroja, K. Gordon, K.G. Millard, V. Ramos, F. Schmidt, Y. Weisblum, D. Jha, M. Tankelevich, G. Martinez-Delgado, J. Yee, R. Patel, J. Dizon, C. Unson-O’Brien, I. Shimeliovich, D.F. Robbiani, Z. Zhao, A. Gazumyan, R.E. Schwartz, T. Hatziioannou, P.J. Bjorkman, S. Mehandru, P.D. Bieniasz, M. Caskey, and M.C. Nussenzweig. 2021b. Evolution of antibody immunity to SARS-CoV-2. Nature 591:639–644.

Goel, R.R., M.M. Painter, K.A. Lundgreen, S.A. Apostolidis, A.E. Baxter, J.R. Giles, D. Mathew, A. Pattekar, A. Reynaldi, D.S. Khoury, S. Gouma, P. Hicks, S. Dysinger, A. Hicks, H. Sharma, S. Herring, S. Korte, W. Kc, D.A. Oldridge, R.I. Erickson, M.E. Weirick, C.M. McAllister, M. Awofolaju, N. Tanenbaum, J. Dougherty, S. Long, K. D’Andrea, J.T. Hamilton, M. McLaughlin, J.C. Williams, S. Adamski, O. Kuthuru, E.M. Drapeau, M.P. Davenport, S.E. Hensley, P. Bates, A.R. Greenplate, and E.J. Wherry. 2022. Efficient recall of Omicron-reactive B cell memory after a third dose of SARS-CoV-2 mRNA vaccine. Cell 185:1875–1887 e1878.

Grifoni., A. D. Weiskopf., and S.I. Ramirez. 2020. Targets of T cell responses to SARS-CoV-2 coronavirus in humans with COVID-19 disease and unexposed individuals. Cell

Guo, Y., K. Chen, P.D. Kwong, L. Shapiro, and Z. Sheng. 2019. cAb-Rep: a database of curated antibody repertoires for exploring antibody diversity and predicting antibody prevalence. Frontiers in immunology 2365.

Gupta, N.T., J.A. Vander Heiden, M. Uduman, D. Gadala-Maria, G. Yaari, and S.H. Kleinstein. 2015. Change-O: a toolkit for analyzing large-scale B cell immunoglobulin repertoire sequencing data. Bioinformatics 31:3356–3358.

Hachmann, N.P., J. Miller, A.-r.Y. Collier, J.D. Ventura, J. Yu, M. Rowe, E.A. Bondzie, O. Powers, N. Surve, K. Hall, and D.H. Barouch. 2022. Neutralization Escape by the SARS-CoV-2 Omicron Variants BA.2.12.1 and BA.4/BA.5. medRxiv 2022.2005.2016.22275151.

Kaku, C.I., A.J. Bergeron, C. Ahlm, J. Normark, M. Sakharkar, M.N.E. Forsell, and L.M. Walker. 2022. Recall of pre-existing cross-reactive B cell memory following Omicron BA.1 breakthrough infection. Sci Immunol eabq3511.

Kuhlmann, C., C.K. Mayer, M. Claassen, T. Maponga, W.A. Burgers, R. Keeton, C. Riou, A.D. Sutherland, T. Suliman, M.L. Shaw, and W. Preiser. 2022. Breakthrough infections with SARS-CoV-2 omicron despite mRNA vaccine booster dose. Lancet 399:625–626.

Liu, C., H.M. Ginn, W. Dejnirattisai, P. Supasa, B. Wang, A. Tuekprakhon, R. Nutalai, D. Zhou, A.J. Mentzer, Y. Zhao, H.M.E. Duyvesteyn, C. Lopez-Camacho, J. Slon-Campos, T.S. Walter, D. Skelly, S.A. Johnson, T.G. Ritter, C. Mason, S.A. Costa Clemens, F. Gomes Naveca, V. Nascimento, F. Nascimento, C. Fernandes da Costa, P.C. Resende, A. Pauvolid-Correa, M.M. Siqueira, C. Dold, N. Temperton, T. Dong, A.J. Pollard, J.C. Knight, D. Crook, T. Lambe, E. Clutterbuck, S. Bibi, A. Flaxman, M. Bittaye, S. Belij-Rammerstorfer, S.C. Gilbert, T. Malik, M.W. Carroll, P. Klenerman, E. Barnes, S.J. Dunachie, V. Baillie, N. Serafin, Z. Ditse, K. Da Silva, N.G. Paterson, M.A. Williams, D.R. Hall, S. Madhi, M.C. Nunes, P. Goulder, E.E. Fry, J. Mongkolsapaya, J. Ren, D.I. Stuart, and G.R. Screaton. 2021. Reduced neutralization of SARS-CoV-2 B.1.617 by vaccine and convalescent serum. Cell 184:4220–4236 e4213.

Madhi, S.A., G. Kwatra, J.E. Myers, W. Jassat, N. Dhar, C.K. Mukendi, A.J. Nana, L. Blumberg, R. Welch, N. Ngorima-Mabhena, and P.C. Mutevedzi. 2022. Population Immunity and Covid-19 Severity with Omicron Variant in South Africa. N Engl J Med 386:1314–1326.

Mallapaty, S. 2022. COVID-19: How Omicron overtook Delta in three charts. Nature

Muecksch, F., Z. Wang, A. Cho, C. Gaebler, T. Ben Tanfous, J. DaSilva, E. Bednarski, V. Ramos, S. Zong, B. Johnson, R. Raspe, D. Schaefer-Babajew, I. Shimeliovich, M. Daga, K.H. Yao, F. Schmidt, K.G. Millard, M. Turroja, M. Jankovic, T.Y. Oliveira, A. Gazumyan, M. Caskey, T. Hatziioannou, P.D. Bieniasz, and M.C. Nussenzweig. 2022. Increased Memory B Cell Potency and Breadth After a SARS-CoV-2 mRNA Boost. Nature

Muecksch, F., Y. Weisblum, C.O. Barnes, F. Schmidt, D. Schaefer-Babajew, Z. Wang, J.C. Lorenzi, A.I. Flyak, A.T. DeLaitsch, and K.E. Huey-Tubman. 2021. Affinity maturation of SARS-CoV-2 neutralizing antibodies confers potency, breadth, and resilience to viral escape mutations. Immunity 54:1853-1868. e1857.

Nealon, J., and B.J. Cowling. 2022. Omicron severity: milder but not mild. Lancet 399:412–413.

Nemet, I., L. Kliker, Y. Lustig, N. Zuckerman, O. Erster, C. Cohen, Y. Kreiss, S. Alroy-Preis, G. Regev-Yochay, E. Mendelson, and M. Mandelboim. 2022. Third BNT162b2 Vaccination Neutralization of SARS-CoV-2 Omicron Infection. N Engl J Med 386:492–494.

Nutalai, R., D. Zhou, A. Tuekprakhon, H.M. Ginn, P. Supasa, C. Liu, J. Huo, A.J. Mentzer, H.M.E. Duyvesteyn, A. Dijokaite-Guraliuc, D. Skelly, T.G. Ritter, A. Amini, S. Bibi, S. Adele, S.A. Johnson, B. Constantinides, H. Webster, N. Temperton, P. Klenerman, E. Barnes, S.J. Dunachie, D. Crook, A.J. Pollard, T. Lambe, P. Goulder, C. Conlon, A. Deeks, J. Frater, L. Frending, S. Gardiner, A. Jämsén, K. Jeffery, T. Malone, E. Phillips, L. Rothwell, L. Stafford, N.G. Paterson, M.A. Williams, D.R. Hall, J. Mongkolsapaya, E.E. Fry, W. Dejnirattisai, J. Ren, D.I. Stuart, and G.R. Screaton. 2022. Potent cross-reactive antibodies following Omicron breakthrough in vaccinees. Cell

Park, Y.-J., D. Pinto, A.C. Walls, Z. Liu, A.D. Marco, F. Benigni, F. Zatta, C. Silacci-Fregni, J. Bassi, K.R. Sprouse, A. Addetia, J.E. Bowen, C. Stewart, M. Giurdanella, C. Saliba, B. Guarino, M.A. Schmid, N. Franko, J. Logue, H.V. Dang, K. Hauser, J. di Iulio, W. Rivera, G. Schnell, F.A. Lempp, J. Janer, R. Abdelnabi, P. Maes, P. Ferrari, A. Ceschi, O. Giannini, G. Dias de Melo, L. Kergoat, H. Bourhy, J. Neyts, L. Soriaga, L.A. Purcell, G. Snell, S.P.J. Whelan, A. Lanzavecchia, H.W. Virgin, L. Piccoli, H. Chu, M.S. Pizzuto, D. Corti, and D. Veesler. 2022. Imprinted antibody responses against SARS-CoV-2 Omicron sublineages. bioRxiv 2022.2005.2008.491108.

Quandt, J., A. Muik, N. Salisch, B.G. Lui, S. Lutz, K. Kruger, A.K. Wallisch, P. Adams-Quack, M. Bacher, A. Finlayson, O. Ozhelvaci, I. Vogler, K. Grikscheit, S. Hoehl, U. Goetsch, S. Ciesek, O. Tureci, and U. Sahin. 2022. Omicron BA.1 breakthrough infection drives cross-variant neutralization and memory B cell formation against conserved epitopes. Sci Immunol eabq2427.

Richardson, S.I., V.S. Madzorera, H. Spencer, N.P. Manamela, M.A. van der Mescht, B.E. Lambson, B. Oosthuysen, F. Ayres, Z. Makhado, T. Moyo-Gwete, N. Mzindle, T. Motlou, A. Strydom, A. Mendes, H. Tegally, Z. de Beer, T. Roma de Villiers, A. Bodenstein, G. van den Berg, M. Venter, T. de Oliviera, V. Ueckermann, T.M. Rossouw, M.T. Boswell, and P.L. Moore. 2022. SARS-CoV-2 Omicron triggers cross-reactive neutralization and Fc effector functions in previously vaccinated, but not unvaccinated, individuals. Cell Host Microbe

Robbiani, D.F., C. Gaebler, F. Muecksch, J.C.C. Lorenzi, Z. Wang, A. Cho, M. Agudelo, C.O. Barnes, A. Gazumyan, S. Finkin, T. Hagglof, T.Y. Oliveira, C. Viant, A. Hurley, H.H. Hoffmann, K.G. Millard, R.G. Kost, M. Cipolla, K. Gordon, F. Bianchini, S.T. Chen, V. Ramos, R. Patel, J. Dizon, I. Shimeliovich, P. Mendoza, H. Hartweger, L. Nogueira, M. Pack, J. Horowitz, F. Schmidt, Y. Weisblum, E. Michailidis, A.W. Ashbrook, E. Waltari, J.E. Pak, K.E. Huey-Tubman, N. Koranda, P.R. Hoffman, A.P. West, Jr., C.M. Rice, T. Hatziioannou, P.J. Bjorkman, P.D. Bieniasz, M. Caskey, and M.C. Nussenzweig. 2020. Convergent antibody responses to SARS-CoV-2 in convalescent individuals. Nature 584:437–442.

Schmidt, F., F. Muecksch, Y. Weisblum, J. Da Silva, E. Bednarski, A. Cho, Z. Wang, C. Gaebler, M. Caskey, and M.C. Nussenzweig. 2022. Plasma neutralization of the SARS-CoV-2 Omicron variant. New Engl J Med 386:599–601.

Schmidt, F., Y. Weisblum, F. Muecksch, H.-H. Hoffmann, E. Michailidis, J.C. Lorenzi, P. Mendoza, M. Rutkowska, E. Bednarski, and C. Gaebler. 2020a. Measuring SARS-CoV-2 neutralizing antibody activity using pseudotyped and chimeric viruses. Journal of Experimental Medicine 217:

Schmidt, F., Y. Weisblum, F. Muecksch, H.H. Hoffmann, E. Michailidis, J.C.C. Lorenzi, P. Mendoza, M. Rutkowska, E. Bednarski, C. Gaebler, M. Agudelo, A. Cho, Z. Wang, A. Gazumyan, M. Cipolla, M. Caskey, D.F. Robbiani, M.C. Nussenzweig, C.M. Rice, T. Hatziioannou, and P.D. Bieniasz. 2020b. Measuring SARS-CoV-2 neutralizing antibody activity using pseudotyped and chimeric viruses. J Exp Med 217:

Seaman, M.S., M.J. Siedner, J. Boucau, C.L. Lavine, F. Ghantous, M.Y. Liew, J. Mathews, A. Singh, C. Marino, J. Regan, R. Uddin, M.C. Choudhary, J.P. Flynn, G. Chen, A.M. Stuckwisch, T. Lipiner, A. Kittilson, M. Melberg, R.F. Gilbert, Z. Reynolds, S.L. Iyer, G.C. Chamberlin, T.D. Vyas, J.M. Vyas, M.B. Goldberg, J. Luban, J.Z. Li, A.K. Barczak, and J.E. Lemieux. 2022. Vaccine Breakthrough Infection with the SARS-CoV-2 Delta or Omicron (BA.1) Variant Leads to Distinct Profiles of Neutralizing Antibody Responses. medRxiv

Servellita, V., A.M. Syed, M.K. Morris, N. Brazer, P. Saldhi, M. Garcia-Knight, B. Sreekumar, M.M. Khalid, A. Ciling, P.Y. Chen, G.R. Kumar, A.S. Gliwa, J. Nguyen, A. Sotomayor-Gonzalez, Y. Zhang, E. Frias, J. Prostko, J. Hackett, Jr., R. Andino, D.A. Wadford, C. Hanson, J. Doudna, M. Ott, and C.Y. Chiu. 2022. Neutralizing immunity in vaccine breakthrough infections from the SARS-CoV-2 Omicron and Delta variants. Cell 185:1539–1548 e1535.

Sokal, A., P. Chappert, G. Barba-Spaeth, A. Roeser, S. Fourati, I. Azzaoui, A. Vandenberghe, I. Fernandez, A. Meola, M. Bouvier-Alias, E. Crickx, A. Beldi-Ferchiou, S. Hue, L. Languille, M. Michel, S. Baloul, F. Noizat-Pirenne, M. Luka, J. Megret, M. Menager, J.M. Pawlotsky, S. Fillatreau, F.A. Rey, J.C. Weill, C.A. Reynaud, and M. Mahevas. 2021. Maturation and persistence of the anti-SARS-CoV-2 memory B cell response. Cell 184:1201–1213 e1214.

Soto, C., R.G. Bombardi, A. Branchizio, N. Kose, P. Matta, A.M. Sevy, R.S. Sinkovits, P. Gilchuk, J.A. Finn, and J.E. Crowe. 2019. High frequency of shared clonotypes in human B cell receptor repertoires. Nature 566:398–402.

Supasa, P., D. Zhou, W. Dejnirattisai, C. Liu, A.J. Mentzer, H.M. Ginn, Y. Zhao, H.M.E. Duyvesteyn, R. Nutalai, A. Tuekprakhon, B. Wang, G.C. Paesen, J. Slon-Campos, C. Lopez-Camacho, B. Hallis, N. Coombes, K.R. Bewley, S. Charlton, T.S. Walter, E. Barnes, S.J. Dunachie, D. Skelly, S.F. Lumley, N. Baker, I. Shaik, H.E. Humphries, K. Godwin, N. Gent, A. Sienkiewicz, C. Dold, R. Levin, T. Dong, A.J. Pollard, J.C. Knight, P. Klenerman, D. Crook, T. Lambe, E. Clutterbuck, S. Bibi, A. Flaxman, M. Bittaye, S. Belij-Rammerstorfer, S. Gilbert, D.R. Hall, M.A. Williams, N.G. Paterson, W. James, M.W. Carroll, E.E. Fry, J. Mongkolsapaya, J. Ren, D.I. Stuart, and G.R. Screaton. 2021. Reduced neutralization of SARS-CoV-2 B.1.1.7 variant by convalescent and vaccine sera. Cell 184:2201–2211 e2207.

Victora, G.D., and M.C. Nussenzweig. 2022. Germinal centers. Annual review of immunology 40:413–442.

Wang, Z., J.C. Lorenzi, F. Muecksch, S. Finkin, C. Viant, C. Gaebler, M. Cipolla, H.-H. Hoffmann, T.Y. Oliveira, and D.A. Oren. 2021a. Enhanced SARS-CoV-2 neutralization by dimeric IgA. Science translational medicine 13:eabf1555.

Wang, Z., F. Muecksch, D. Schaefer-Babajew, S. Finkin, C. Viant, C. Gaebler, H.-H. Hoffmann, C.O. Barnes, M. Cipolla, and V. Ramos. 2021b. Naturally enhanced neutralizing breadth against SARS-CoV-2 one year after infection. Nature 595:426–431.

Wang, Z., F. Muecksch, D. Schaefer-Babajew, S. Finkin, C. Viant, C. Gaebler, H.H. Hoffmann, C.O. Barnes, M. Cipolla, V. Ramos, T.Y. Oliveira, A. Cho, F. Schmidt, J. Da Silva, E. Bednarski, L. Aguado, J. Yee, M. Daga, M. Turroja, K.G. Millard, M. Jankovic, A. Gazumyan, Z. Zhao, C.M. Rice, P.D. Bieniasz, M. Caskey, T. Hatziioannou, and M.C. Nussenzweig. 2021c. Naturally enhanced neutralizing breadth against SARS-CoV-2 one year after infection. Nature 595:426–431.

Wang, Z., F. Schmidt, Y. Weisblum, F. Muecksch, C.O. Barnes, S. Finkin, D. Schaefer-Babajew, M. Cipolla, C. Gaebler, J.A. Lieberman, T.Y. Oliveira, Z. Yang, M.E. Abernathy, K.E. Huey-Tubman, A. Hurley, M. Turroja, K.A. West, K. Gordon, K.G. Millard, V. Ramos, J. Da Silva, J. Xu, R.A. Colbert, R. Patel, J. Dizon, C. Unson-O’Brien, I. Shimeliovich, A. Gazumyan, M. Caskey, P.J. Bjorkman, R. Casellas, T. Hatziioannou, P.D. Bieniasz, and M.C. Nussenzweig. 2021d. mRNA vaccine-elicited antibodies to SARS-CoV-2 and circulating variants. Nature 592:616–622.

Weisblum, Y., F. Schmidt, F. Zhang, J. DaSilva, D. Poston, J.C. Lorenzi, F. Muecksch, M. Rutkowska, H.-H. Hoffmann, and E. Michailidis. 2020. Escape from neutralizing antibodies by SARS-CoV-2 spike protein variants. eLife 9:e61312.

Whitaker, M., J. Elliott, B. Bodinier, W. Barclay, H. Ward, G. Cooke, C.A. Donnelly, M. Chadeau-Hyam, and P. Elliott. 2022. Variant-specific symptoms of COVID-19 among 1,542,510 people in England. medRxiv 2022.2005.2021.22275368.

Wolter, N., W. Jassat, S. Walaza, R. Welch, H. Moultrie, M. Groome, D.G. Amoako, J. Everatt, J.N. Bhiman, C. Scheepers, N. Tebeila, N. Chiwandire, M. du Plessis, N. Govender, A. Ismail, A. Glass, K. Mlisana, W. Stevens, F.K. Treurnicht, Z. Makatini, N.Y. Hsiao, R. Parboosing, J. Wadula, H. Hussey, M.A. Davies, A. Boulle, A. von Gottberg, and C. Cohen. 2022. Early assessment of the clinical severity of the SARS-CoV-2 omicron variant in South Africa: a data linkage study. Lancet 399:437–446.

World Health, O. 2022. WHO SAGE roadmap for prioritizing uses of COVID-19 vaccines: an approach to optimize the global impact of COVID-19 vaccines, based on public health goals, global and national equity, and vaccine access and coverage scenarios, first issued 20 October 2020, updated: 13 November 2020, updated: 16 July 2021, latest update: 21 January 2022. In World Health Organization, Geneva.

Yuan, M., H. Liu, N.C. Wu, C.-C.D. Lee, X. Zhu, F. Zhao, D. Huang, W. Yu, Y. Hua, and H. Tien. 2020. Structural basis of a shared antibody response to SARS-CoV-2. Science 369:1119–1123.

